# Improving the accuracy of single turnover active fluorometry (STAF) for the estimation of phytoplankton primary productivity (PhytoPP)

**DOI:** 10.1101/583591

**Authors:** Tobias G. Boatman, Richard J. Geider, Kevin Oxborough

## Abstract

Photosystem II (PSII) photochemistry is the ultimate source of reducing power for phytoplankton primary productivity (PhytoPP). Single turnover active chlorophyll fluorometry (STAF) provides a non-intrusive method that has the potential to measure PhytoPP on much wider spatiotemporal scales than is possible with more direct methods such as ^14^C fixation and O_2_ evolved through water oxidation. Application of a STAF-derived absorption coefficient for PSII light-harvesting (a_LHII_) provides a method for estimating PSII photochemical flux on a unit volume basis (JV_PII_). Within this study, we assess potential errors in the calculation of JV_PII_ arising from sources other than photochemically active PSII complexes (baseline fluorescence) and the package effect. Although our data show that such errors can be significant, we identify fluorescence-based correction procedures that can be used to minimize their impact. For baseline fluorescence, the correction incorporates an assumed consensus PSII photochemical efficiency for dark-adapted material. The error generated by the package effect can be minimized through the ratio of variable fluorescence measured within narrow wavebands centered at 730 nm, where the re-absorption of PSII fluorescence emission is minimal, and at 680 nm, where re-absorption of PSII fluorescence emission is maximal. We conclude that, with incorporation of these corrective steps, STAF can provide a reliable estimate of JV_PII_ and, if used in conjunction with simultaneous satellite measurements of ocean color, could take us significantly closer to achieving the objective of obtaining reliable autonomous estimates of PhytoPP.

## 1 Introduction

Phytoplankton contribute approximately half the photosynthesis on the planet (Field, 1998), thus forming the base of marine food webs. Reliable assessment of Phytoplankton Primary Productivity (PhytoPP) is crucial to an understanding of the global carbon and oxygen cycles and oceanic ecosystem function. Consequently, PhytoPP has been recognized as an Essential Ocean Variable (EOV) within the Global Ocean Observing System (GOOS). PhytoPP is a dynamic biological process that responds to variability in multiple environmental drivers including light, temperature and nutrients across spatial scales from meters to ocean basins, and time scales from minutes to tens of years. This poses significant challenges for measuring and monitoring PhytoPP.

Historically, the most frequently employed method for assessing PhytoPP has been the fixation of ^14^C within closed systems over several hours of incubation (Marra, 2002; Milligan et al. 2015). Despite the widespread use of the ^14^C method, which has led to measurements of PhytoPP by the ^14^C method providing the database against which remote sensing estimates of primary production are calibrated (Bouman et al. 2018), there is considerable uncertainty in what exactly the ^14^C method measures and the accuracy of bottle-incubation based methods for obtaining PhytoPP in oligotrophic ocean waters (Quay et al. 2010).

According to Marra (2002), the ^14^C technique measures somewhere between net and gross carbon fixation, depending on the length of the incubation. In this context, net carbon fixation is defined as gross carbon fixation minus carbon respiratory losses and light-dependent losses due to photorespiration and light-enhanced mitochondrial respiration (Milligan et al. 2015). Although it may seem intuitive that short incubations should provide a good estimate of gross carbon fixation (and closely match PhytoPP), several authors have reported that short-term ^14^C fixation does not reliably measure net or gross production (e.g. Halsey et al. 2013; Milligan et al. 2015). It should also be noted that short-term, in the context of ^14^C fixation, is several hours incubation. This clearly imposes major limitations on the spatiotemporal scales at which PhytoPP can be assessed using this method.

Gross photosynthesis by phytoplankton is defined here as the rate at which reducing power is generated by photosystem II (PSII) through the conversion of absorbed light energy (PSII photochemistry). Within this study, gross photosynthesis is quantified by measuring the rate at which O_2_ is evolved through water oxidation by PSII photochemistry (Ferron et al. 2016) and is termed PhytoGO. Although measurement of O_2_ evolution provides some advantages over ^14^C fixation, in that both gross and net primary production can be obtained, the spatiotemporal limitations are similar.

It is now widely accepted that active fluorometry can provide a non-intrusive method for measuring PSII photochemistry on much wider spatiotemporal scales than either ^14^C fixation or O_2_ evolution. Within oceanic systems, where optically thin conditions are the norm, the most appropriate form of active fluorometry is the single turnover method (Kolber and Falkowski 1993; Kolber et al. 1998; Moore et al. 2006; Suggett et al. 2001; Oxborough et al. 2012). One important parameter generated by single turnover active fluorometry (STAF) is the absorption cross section of PSII photochemistry (σ_PII_ in the dark-adapted state, σ_PII_’ in the light-adapted state, see Terminology) with units of m^2^ PSII^−1^ (Kolber et al. 1998; Oxborough et al. 2012). This parameter allows for the calculation of PSII photochemical flux through a single PSII center, as the product of σ _PII_’ and incident photon irradiance (E, with units of photons m^−2^ s^−1^). PSII photochemical flux has units of photons PSII^−1^ s^−1^ or (assuming an efficiency of one stable photochemical event per photon) electrons PSII^−1^ s^−1^ (Equation 1).

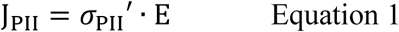

Both PhytoPP and PhytoGO can be reported per unit volume (SI units of C m^−3^ s^−1^ or O_2_ m^−3^ s^−1^, respectively). Given that J_PII_ provides the photochemical flux through the σ_PII_’ provided by a single PSII, the PSII photochemical flux per unit volume (JV_PII_, with units of electrons m^−3^ s^−1^) can be defined as the flux through the absorption cross section of PSII photochemistry provided by all open PSII centers within the volume (Equation 2).

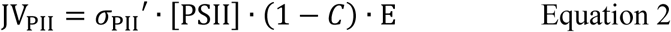

Where [PSII] is the concentration of photochemically active PSII complexes, with units of PSII m^−3^, and (1 - *C*) is the proportion of these centers that are in the open state at the point of measurement under actinic light. It follows that JV_PII_ can, in principle, provide a proxy for PhytoPP (Oxborough et al. 2012).

An important caveat to using JV_PII_ as a proxy for PhytoPP is that there are a number of processes operating within phytoplankton that can uncouple PhytoPP from PhytoGO and PhytoGO from PSII photochemistry (Geider and MacIntyre 2002; Behrenfeld et al. 2004; Halsey et al. 2010; Suggett et al. 2010; Lawrenz et al. 2013). It follows that JV_PII_ provides an upper limit for PhytoPP which is defined by the release of each O_2_ requiring a minimum of four photochemical events and each O_2_ released resulting in the maximum assimilation of one CO_2_.

Previous studies obtained a value for the [PSII] term within Equation 2 from discrete samples of chlorophyll *a* by assuming that the number of PSII centers per chlorophyll *a* (*n*_PSII_) is relatively constant (Kolber & Falkowski, 1993; Suggett et al., 2001). A significant problem with this approach is that *n*_PSII_ shows significant variability, both in laboratory-based cultures (Suggett et al. 2004) and in natural phytoplankton communities (Moore et al. 2006; Suggett et al. 2006). In addition, the derivation of nPSII requires a chlorophyll *a* extraction for each sample: a requirement that imposes significant spatiotemporal limitations.

A STAF-based method for the determination of [PSII] was described by Oxborough at al. (2012). This method operates on the assumption that the ratio of rate constants for PSII photochemistry (*k*_PII_) and PSII fluorescence emission (*k*_FII_) falls within a narrow range across all phytoplankton types. One consequence of this assumption is illustrated by Equation 3.

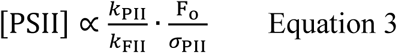

Where F_o_ is the ‘origin’ of variable fluorescence from a dark-adapted sample (see Terminology). Data from a follow-up study (Silsbe et al. 2015), were entirely consistent with Equation 3 and were used to derive a sensor type-specific constant, termed K_a_, for the FastOcean fluorometer (CTG Ltd, West Molesey, UK). It follows that:

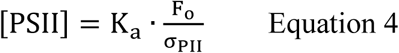

It is worth noting that K_a_ and [PSII], within Equation 4, are spectrally independent while, for a homogeneous population, F_o_ and σ_PII_ are expected to covary with measurement LED intensity.

As noted by Oxborough et al. (2012), Equation 3 is only valid if a high proportion of the fluorescence signal at F_o_ comes from PSII complexes that are photochemically active and in the open state. While it is reasonable to expect that most photochemically active PSII complexes will be in the open state at *t* = 0 during a STAF measurement, there are situations where a significant proportion of the fluorescence signal at F_o_ may come from sources other than photochemically active PSII complexes. This becomes a concern when the observed ratio of variable fluorescence (F_v_) to maximum fluorescence (F_m_) from a dark-adapted sample is low: although the maximum observed F_v_/F_m_ varies among phytoplankton taxa, it is generally within the range of 0.5 to 0.6 for the fluorometers used within this study.

One plausible explanation for sub-maximal F_v_/F_m_ values is that PSII photochemistry is downregulated by high levels of Stern-Volmer quencher within the PSII pigment matrix. As with measurement LED intensity, F_o_ and σ_PII_ covary with Stern-Volmer quenching and Equation 4 remains valid. Light-dependent accumulation of Stern-Volmer quencher within the PSII pigment matrix generates non-photochemical quenching of PSII fluorescence (NPQ) within a wide range of phytoplankton groups (Olaizola and Yamamoto 1994; Demmig-Adams and Adams 2006; Goss and Jakob 2010; Krause and Jahns, 2004). However, this form of quenching is generally reversed within tens of seconds to a few minutes dark-adaptation and would therefore not be expected to significantly decrease F_v_/F_m_.

A second plausible explanation for sub-maximal F_v_/F_m_ values is that a proportion of the signal at F_o_ is generated by PSII complexes that lack photochemically active reaction centers. Under the assumption that these complexes are not energetically coupled to photochemically active PSII complexes, their presence would increase Fo but have no impact on s_PII_. Consequently, the value of [PSII] generated by Equation 4 would increase in proportion to the increase in measured F_o_. Within this manuscript, the fraction of F_o_ that does not originate from open PSII complexes is termed baseline fluorescence (F_b_) and the fraction that does is termed baseline-corrected F_o_ (F_oc_, see Terminology).

Within Equation 2, JV_PII_ is proportional to the product of [PSII] and (1 − *C*) during a STAF measurement under actinic light. A value for the concentration of photochemically active PSII centers can be generated from a STAF measurement made on a dark-adapted sample using Equation 4. The proportion of these complexes in the open state has routinely been estimated through the qP parameter (Kolber et al. 1998) which is mathematically equivalent to the photochemical factor (F_q_’/F_v_’) defined by Baker and Oxborough (2004). This requires determination of F_o_’, using the equation provided by Oxborough and Baker (1997) or through direct measurement after 1 – 2 s dark-adaptation following a STAF measurement under actinic light (Kolber et al. 1998).

As an alternative to Equation 2, Oxborough et al. (2012) include a method for calculating JV_PII_ that does not require [PSII], (I – *C*) or σ_PII_ (Equation 5).

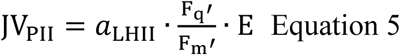

Where a_LHII_ is the absorption coefficient for PSII light harvesting, with SI units of m^−1^. A value for a_LHII_ can be derived using Equation 6.

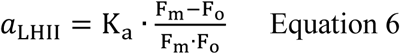

The link between Equations 2 and 5 is illustrated by Equation 7 (Kolber et al. 1998) and Equation 8 (Oxborough et al. 2012).

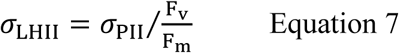

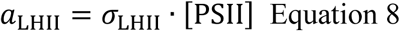

The package effect is a consequence of the high concentration of chlorophyll a and other light-absorbing pigments within phytoplankton cells. To put this in context, while the concentration of chlorophyll a within the open ocean is often below 0.1 mg m^−3^, the concentration within phytoplankton cells is approximately a million times higher than this, at 0.1 kg m^−3^ (calculated from data within Montagnes et al. 1994). It follows that while sea water with phytoplankton cells suspended within it can be considered optically thin, the localized volume within each phytoplankton cell is optically very thick.

Differences in the package effect due to pigment composition and morphology among species have been identified (Berner et al., 1989; Bricaud et al. 1993; Morel and Bricaud, 1981). Even within individual phytoplankton species, levels of pigment packing vary with eco-physiological condition and life cycle (Berner et al. 1989; Falkowski and LaRoche, 1991). Increases in the magnitude of the package effect will increase the absorption of photons generated by PSII fluorescence (FII) before these photons leave the cell, and thus decrease the measured value of FII relative to PSII photochemistry (PII). Given that a fundamental assumption of the absorption method is that the relationship between PSII photochemistry (PII) and PSII fluorescence emission (FII) is reasonably constant, variability in the level of package effect among samples clearly has the potential to introduce significant errors.

The main objective for this study was to test the applicability of Equations 4 and 6. Because the values generated by both equations are dependent on K_a_, a comprehensive evaluation of the absolute value and general applicability of K_a_ has been incorporated within the study. As a first step, a large number of sample-specific values of K_a_ were generated by combining data from parallel STAF and flash O_2_ measurements from eleven phytoplankton species, grown under nutrient-replete and N-limited conditions. This allowed an evaluation of the degree to which sub-maximal values of F_v_/F_m_ could be attributed to Stern-Volmer quenching or baseline fluorescence. It should be noted that the term sample-specific K_a_ is used to define apparent K_a_ values that are not corrected for baseline fluorescence. Each sample-specific K_a_ value referenced is the mean of all reps for a specific combination of species and growth conditions (nutrient replete or N-starved).

In addition to the STAF and flash O_2_ measurements used to generate sample-specific values of K_a_, the same measurement systems were used to run fluorescence light curves (FLCs) and oxygen light curves (OLCs) on all eleven phytoplankton species. The data generated from these measurements allowed for direct comparison of PhytoGO and JV_PII_ (from Equation 6) at multiple points through the light curves.

This first set of experiments provided evidence for a wider range of K_a_ values across species and environmental conditions than was evident in the earlier studies of Oxborough et al. (2012) and Silsbe et al. (2015). Although the intra-species variance of K_a_ values (between values determined for nutrient-replete and N-limited cultures) could confidently be linked to baseline fluorescence, the inter-species variance was more easily explained in terms of the package effect. To test this hypothesis, an additional set of measurements were made on 11 phytoplankton species, of which six were common to the first set of experiments. The range of species was selected to cover a wide range of cell sizes and optical characteristics. As before, sample-specific K_a_ values were generated from parallel flash O_2_ and STAF measurements. The STAF measurements were made using FastBallast fluorometers (CTG Ltd, as before) fitted with narrow bandpass filters centered at 680 nm and 730 nm. These wavebands were chosen because chlorophyll a fluorescence is absorbed much more strongly at 680 nm than at 730 nm. It follows that attenuation of fluorescence emission due to the package effect will be much higher at 680 nm than at 730 nm and thus that variability of the package effect among species should correlate with the ratio of fluorescence outputs measured at 730 nm and 680 nm. To allow for comparison with the existing FastOcean sensor, a third FastBallast sensor was fitted with the bandpass filter used within FastOcean.

## 2 Materials and methods

### 2.1 Phytoplankton cultures (N-limited experiments)

Semi-continuous phytoplankton cultures were maintained and adapted to nutrient-replete conditions. All cultures were grown in *f*/2 medium with silicates omitted where appropriate (Guillard, 1975).

The experimental work covered a period of several months. The initial work was conducted at the University of Essex and incorporated six phytoplankton species (Table 1). Cultures were maintained at 20 °C in a growth room (Sanyo Gallenkamp PLC, UK) and illuminated by horizontal fluorescent tubes (Sylvania Luxline Plus FHQ49/T5/840, UK). The Light:Dark (L:D) cycle was set at 12 h:12 h. Neutral density filters were used to generate low light and high light conditions (photon irradiances of 30 and 300 µmol photons m^−2^ s^−1^, respectively). 300 mL culture volumes were maintained within 1 L Duran bottles. Cultures were constantly aerated with ambient air and mixed using magnetic stirrers.

**Table 1.** List of cultures used within each experiment. H = High Light, L = Low Light, N = N-limited. UoE = University of Essex. (* = simultaneous N-limited OLC/FLC measurements made on *T. weissflogii* cultures (*n* = 10).

| Algal species<br>(Symbol used within figures) | Clone | Site(s) | Media | F<sub>b</sub> |  |  |  | Package effect |
| --- | --- | --- | --- | --- | --- | --- | --- | --- |
|  |  |  |  | Flash O<sub>2</sub> |  |  | OLC<br>FLC | STAF/<br>flash O<sub>2</sub> |
|  |  |  |  | H | L | N |  |  |
| *Calcidiscus leptoporus*<br>(*C. l*) | RCC1159 | CTG | f/2 | - | - | - | - | 4 |
| *Chlorella vulgaris*<br>(*C. v*) | CCAP211<br>/12 | CTG | BG11 | - | 8 | 4 | 5 | 4 |
| *Coccolithus pelagicus*<br>(*C. p*) | PCC182 | CTG | f/2<br>(+Si) | - | 8 | 4 | 5 | - |
| *Coscinodiscus granii*<br>(*C. g*) | CCAP1013<br>/10 | CTG | f/2<br>(+Si) | - | - | - | - | 4 |
| *Coscinodiscus* sp.<br>(*C. sp.*) | CCAP1013<br>/11 | CTG | f/2<br>(+Si) | - | - | - | - | 4 |
| *Dunaliella salina*<br>(*D. s*) | CCAP19<br>/18 | UoE<br>CTG | f/2 | 8 | 2 | 4 | 5 | - |
| *Dunaliella tertiolecta*<br>(*D. t*) | CCAP1320 | UoE<br>CTG | f/2 | 6 | 2 | 4 | 5 | 4 |
| *Emiliania huxleyi*<br>(*E. h*) | CCMP1516 | UoE<br>CTG | f/2 | 8 | 2 | 4 | 5 | 4 |
| *Isochrysis galbana*<br>(*I. g*) | CCMP1323 | CTG | f/2 | - | 8 | 4 | 5 | 4 |
| *Pseudo-nitzschia fraudulenta*<br>(*P-n. f*) | CCAP1061<br>/46 | CTG | f/2<br>(+Si) | - | - | - | - | 4 |
| *Pycnococcus provasolii*<br>(*P. p*) | CCMP1199 | CTG | f/2 | - | 8 | 4 | 5 | 4 |
| *Phaeodactylum tricornutum*<br>(*P. t*) | CCMP2561 | UoE<br>CTG | f/2<br>(+Si) | 8 | 2 | 4 | 5 | 4 |
| *Thalassiosira pseudonana*<br>(*T. p*) | CCMP1335 | CTG | f/2<br>(+Si) | - | 8 | 4 | 5 | - |
| *Thalassiosira punctigera*<br>(*C. p*) | CCAP1085<br>/19 | UoE<br>CTG | f/2<br>(+Si) | 8 | 2 | 4 | 5 | - |
| *Thalassiosira rotula*<br>(*T. r*) | CCAP1085<br>/20 | CTG | f/2<br>(+Si) | - | - | - | - | 4 |
| *Thalassiosira weissflogii*<br>(*T. w*) | CCMP1051 | UoE/<br>CTG | f/2<br>(+Si) | 8 | 2 | 4 | 5<br>(10*) | - |

**Table 2.**
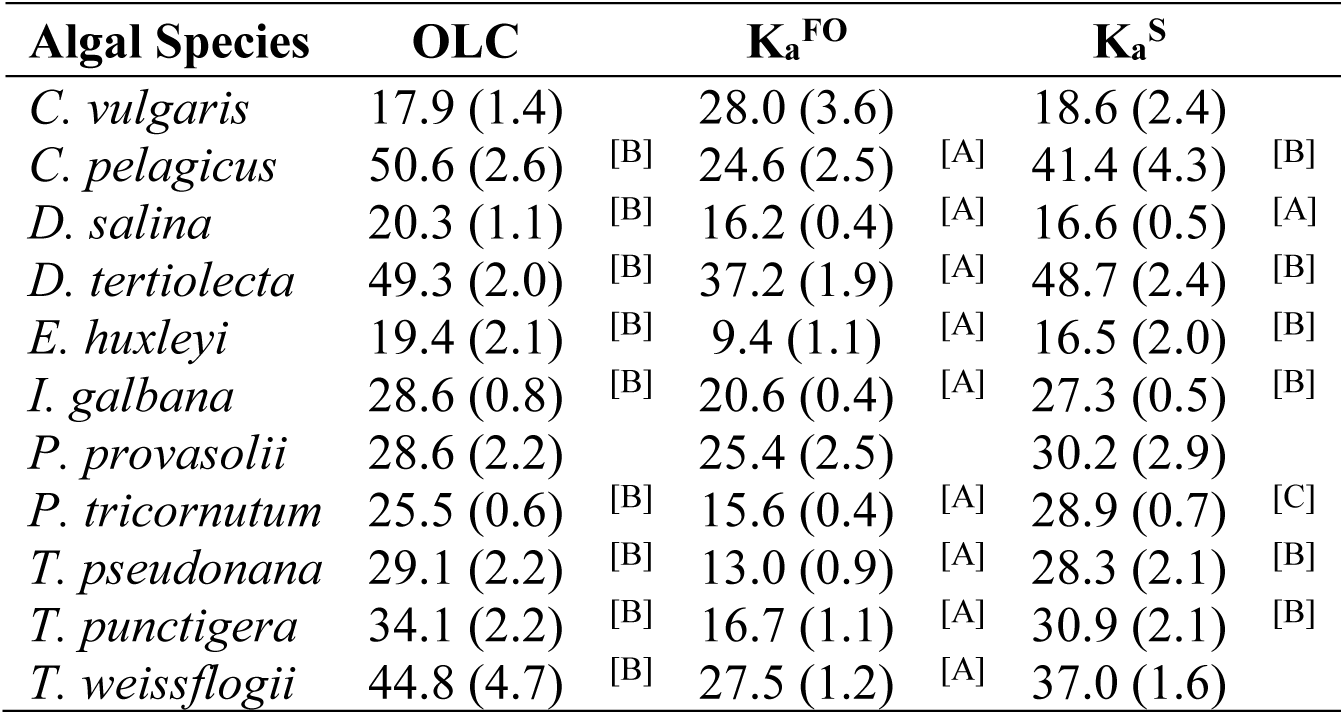
The maximum phytoplankton gross photosynthesis rates (PhytoGO_m_) from simultaneous OLC and FLC measurements of the 11 nutrient-replete phytoplankton cultures measured in Experiment 1. PhytoGO from the FLC data was calculated using K_a_^FO^ (11,800 m^−1^) and a sample-specific (K_a_^S^) values. Differences between OLC and FLC data was tested by a series of parametric One-Way ANOVA tests with a post hoc Tukey test (One-way ANOVA, Tukey post hoc test; P < 0.05). Letters show the significant differences between the maximum gross photosynthesis rate (PhytoGP_m_) (O_2_ RCII^−1^ s^−1^); where [B] is significantly greater than [A], and [C] is significantly greater than [A] and [B].

Additional experiments, incorporating the remaining five species, were conducted at CTG Ltd (as before). Cultures were maintained as 30 mL aliquots within filter-capped tissue culture flasks (Fisher Scientific, UK: 12034917). A growth temperature of 20 °C was maintained by placing the flasks within a water bath (Grant SUB Aqua Pro 2 L, USA). Low light illumination (photon irradiance of 30 µmol photons m^−2^ s^−1^) was provided from white LED arrays (Optoelectronic Manufacturing Corporation Ltd. 1ft T5 Daylight, UK). The L:D cycle was set at 12 h:12 h.

The N-limited cultures were sub-cultured from the nutrient-replete cultures. High light-grown cultures were used for the six species interrogated at the University of Essex. In all cases, the growth photon irradiance of the original culture was maintained after sub-culturing. All N-limited cultures were grown into the stationary growth phase using N-limiting *f*/2 medium before experimental measurements were made.

### 2.2 Phytoplankton cultures (package effect experiments)

All package effect experiments were conducted at CTG Ltd. Cultures were maintained as 30 mL aliquots within filter-capped tissue culture flasks (Fisher Scientific, UK: 12034917). A growth temperature of 20 °C was maintained by placing the flasks within a water bath (Grant SUB Aqua Pro 2 L, USA). Low light illumination (photon irradiance of 30 µmol photons m^−2^ s^−1^) was provided from white LED arrays (Optoelectronic Manufacturing Corporation Ltd. 1ft T5 Daylight, UK). The L:D cycle was set at 12 h:12 h.

### 2.3 Setup for OLCs and flash O_2_ measurements

All OLCs and flash O2 measurements were made using an Oxygraph Plus system (Hansatech Instruments Ltd, Norfolk, UK). The sample volume was always 1.5 mL and a sample temperature of 20 °C was maintained using a circulating water bath connected to the water jacket of the DW1 electrode chamber. The sample was mixed continuously using a magnetic flea (as supplied with the Oxygraph Plus system). Illumination was provided from an Act2 laboratory system (CTG Ltd, as before). The source comprised three blue Act2 LED units incorporated within an Act2 Oxygraph head. Automated control of continuous illumination during OLCs or the delivery of saturating pulses during flash O2 measurements was provided by an Act2 controller and the supplied Act2Run software package.

### 2.4 Dilution of samples between flash O_2_ and STAF measurements

The N-limited and dual waveband experiments included determination of sample-specific K_a_ values. In all cases, the required dark STAF measurements of F_o_ and σ_PII_ were made after the flash O_2_ measurements. In all cases, filtered medium was used to dilute the sample between Oxygraph and STAF measurements.

### 2.5 Chlorophyll *a* extraction

In all cases, the concentrated sample used for flash O_2_ or OLCs was normalized to the parallel dilute STAF sample used to generate F_o_ and σ_PII_ or FLC data through direct measurement of chlorophyll a concentration from both samples.

Chlorophyll was quantified by pipetting 0.5 mL of each sample into 4.5 mL of 90% acetone and extracting overnight in a freezer at −20 °C (Welschmeyer, 1994). Samples were re-suspended and centrifuged at approximately 12,000 × g for 10 minutes and left in the dark (∼ 30 minutes) to equilibrate to ambient temperature. Raw fluorescence from a 2 mL aliquot was measured using a Trilogy laboratory fluorometer (Turner, UK). The chlorophyll a concentration was then calculated from a standard curve.

### 2.6 Setup for dark STAF measurements and FLCs (N-limited experiments)

All STAF measurements for the N-limited experiments were made using a FastOcean sensor in combination with an Act2 laboratory add-on (CTG Ltd, as before). The Act2 FLC head was populated with blue LEDs. A water bath was used as a source for the FLC head water jacket, maintaining the sample temperature at 20 °C.

### 2.7 Flash O_2_ measurements for determining sample-specific K_a_ values

The density of photochemically active PSII complexes within each sample was determined using the flash O_2_ method (Mauzerall & Greenbaum, 1989; Suggett et al., 2003; Silsbe et al. 2015). The standard flash used was 120 µs duration on a 24 ms pitch at a photon irradiance of 22,000 µmol photons m^−2^ s^−1^.

The concentration of photochemically active PSII centers is proportional to the product of gross O_2_ evolution rates (E_0_) and the reciprocal of flash frequency (Hz). The basic theoretical assumptions are that all photochemically active PSII centers undergo stable charge separation once during each flash, that all photochemically active PSII centers re-open before the next flash and that four stable charge separation events are required for each O2 released. In reality, small errors are introduced because some centers do not undergo stable charge separation with each flash (misses) while some centers will undergo more than one stable charge separation event with each flash (multiple hits).

The following checks were applied with all samples:

- The proportion of PSII centers closed during each flash was verified by comparison with sequences of 120 µs flashes on a 24 ms pitch at a photon irradiance of 13,800 µmol photons m^−2^ s^−1^
- The default flash pitch of 24 ms was compared against 16 ms and 36 ms to assess the accumulation of closed PSII centers, with 120 µs flashes of 22,000 µmol photons m^−2^ s^−1^ being applied in all three cases
- Sequences of 180 and 240 µs flashes on a 24 ms pitch at a photon irradiance of 22,000 µmol photons m^−2^ s^−1^ were applied to assess multiple hits

In all cases, a flash duration of 120 µs duration at a photon irradiance of 22,000 µmol photons m^−2^ s^−1^ on a 24 ms pitch provided more than 96% saturation, with no evidence of a significant level of multiple hits or the accumulation of closed PSII centers.

### 2.8 Parallel OLC and FLC measurements (N-limited experiments)

A series of parallel replicate OLC/FLC measurements were made on all nutrient-replete cultures, as well as for the N-limited T. weissflogii culture (Table 1). The 10 to 12 light steps were identical between the parallel OLC and FLC measurements. The sequences always started with a dark step, with all subsequent steps lasting 180 s. Additional dark steps were incorporated after every third light step. The dark respiration rate (Rd) was assessed before, during and after the OLC. The Rd values measured during and after the OLC were always within 8% of the initial Rd (n = 65). The FastOcean ST sequence comprised 100 flashlets on a 2 µs pitch. Each acquisition was an average of 16 sequences on a 100 ms pitch. The auto-LED and auto-PMT functions incorporated within the Act2Run software were always active.

The reported gross O_2_ evolution rates (E_0_) were taken as the sum of measured net O_2_ evolution (P_n_) and R_d_ (Equation 9).

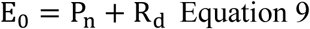

### 2.9 OLC and FLC curve fits (N-limited experiments)

OLCs and FLCs are variants of the widely used P-E (photosynthesis – photon irradiance) curve. For OLCs, the metric for photosynthesis is the rate at which O_2_ is evolved through water oxidation by PSII. For FLCs, the metric for photosynthesis is the relative rate of PSII photochemistry, which is assessed as the product of ϕ_PII_ and E. In the absence of baseline fluorescence (when F_b_ = 0), the parameter F_q_’/F_m_’ can be used to provide an estimate of ϕ_PII_. It follows that FLC curves can be generated by plotting E against the product of baseline-corrected F_q_’/F_m_’ (F_q_’/F_mc_’) and E.

There are three basic parameters derived from all P-E curve fits: α, E_k_ and P_m_. The value of α provides the initial slope of the relationship between E and P. E_k_ is an inflection point along the P-E curve which is often described as the light saturation parameter (Platt and Gallegos, 1980). P_m_ is the maximum rate of photosynthesis.

The FLC curve fits within this study were generated by the Act2Run software (CTG Ltd, as before). The curve fitting routine within Act2Run is a two-step process which takes advantage of the fact that the signal to noise within FLC data is highest during the initial part of the FLC curve. In the first step (the Alpha phase), Equation 10 is used to generate values for α and E_k_ (Webb et al. 1974; Silsbe and Kromkamp, 2012). The overall fit is an iterative process that minimizes the sum of squares of the difference between observed and fit values. During the Alpha fit, a significant weighting on the initial points (low actinic E values) is generated by multiplying each square of the difference by (F_q_’/F_mc_’)2. This approach normally generates a good fit up to E_k_, but overshoots beyond this point. Consequently, the P_m_ values generated by the Alpha phase are generally too high.

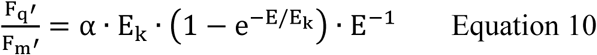

In the second step (the Beta phase) Equation 11 is used to improve the value of P_m_. This step includes a second exponential which is only applied to data points at E values above the E_k_ value generated by the Alpha phase. The sum of squares of the difference between observed and fit values is not weighted during the Beta phase. This approach forces ϕ_PII_ at E_k_ to be 63.2% of α.

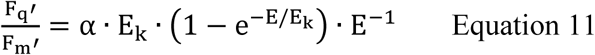

The signal to noise for OLC data tends to increase with E (the opposite of what happens with FLC data). Consequently, the fitting method used for the FLC data is not appropriate for OLC data as it is highly dependent on having a good signal to noise during the early part of the curve. The iterative OLC data fits used Equations 12 and 13 (Platt and Gallegos, 1980).

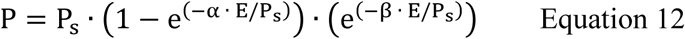

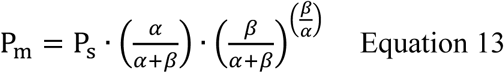

Within these equations, P_s_ and β improve some fits by incorporating a phase that accounts for possible photoinactivation of PSII complexes and/or supra-optimal levels of PSII downregulation (photoinhibition).

Direct comparison of α and β between the FLC and OLC is problematic because while □ is incorporated through the entire curve for both fits, β is only incorporated beyond E_k_ for the FLC and through the entire curve for the OLC. For this reason, direct comparison between FLC and OLC data was focused on P_m_ values.

### 2.10 Setup for the package effect STAF measurements

STAF measurements were made using three FastBallast sensors (CTG Ltd, as before). Each sensor was fitted with one of the following bandpass filters:

730 nm bandpass, 10 nm FWHM (Edmund Optics, UK; part number 65-176)

680 nm bandpass, 10 nm FWHM (Edmund Optics, UK; part number 88-571)

682 nm bandpass, 30 nm FWHM (HORIBA Scientific, UK; part number 682AF30)

Where FWHM is Full Width at Half Maximum. The FastBallast units fitted with filters A, B and C are, hereafter, termed B730, B680 and B682, respectively. Filter C is the standard bandpass filter fitted within FastOcean and FastBallast fluorometers and was included here for comparison.

The emission peak for PSII fluorescence is cantered at 683 nm and is Stokes shifted from a strong absorption peak cantered at 680 nm. Consequently, reabsorption of PSII fluorescence defined by B680 is close to maximal and is also very high when PSII fluorescence is defined by B682. In contrast, reabsorption of PSII fluorescence within the waveband defined by B730 is minimal.

Because the FastBallast sensor does not incorporate a water jacket, all measurements were made in a temperature-controlled room at 20 °C. The FastBallast units were always switched on immediately before each test and automatically powered down once a test had finished. This procedure prevented any measurable increase in temperature within the FastBallast sample chamber during testing.

Calibration of FastBallast units does not include as assessment of ELED (Equation 4). Consequently, there is no instrument-type specific K_a_ available for FastBallast. To get around this limitation, the LED output was maintained at a constant level for all measurements. This allowed Equation 16 to be used in place of Equation 4.

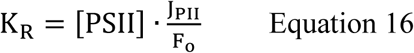

Where K_R_ is the instrument-specific constant defined by Oxborough et al. (2012) with units of photons m^−3^ s^−1^ and JPII is the initial rate of PSII photochemical flux during a STAF pulse, with units of electrons PSII^−1^ s^−1^. As before, it is assumed that each photon used to drive PSII photochemistry results in the transfer of one electron.

Samples used for the FastBallast STAF measurements were prepared by diluting a small aliquot from the sample used for the associated flash O_2_ measurement. 60 mL of cell suspension was prepared and divided equally among the three FastBallast sample chambers. Samples were dark-adapted for at least five minutes before measurements were made.

Each sample was run through the three FastBallast units simultaneously using the FaBtest software supplied with the FastBallast fluorometer (CTG Ltd, as before). This test involves continuous application of 400 µs saturating pulses at 10 Hz to a slowly stirred 20 mL sample for eight minutes. Only 0.5 mL of the sample is illuminated by the saturating pulse at any point in time. Stirring the sample ensures that the entire sample is interrogated during the test and prevents the accumulation of closed PSII reaction centers. The mean value of F_v_ was extracted from each test result.

Following the initial measurement, a spike of Basic Blue 3 (BB3) was added to each sample to increase the extracellular baseline fluorescence. BB3 is a water-soluble fluorescent dye, which absorbs throughout the visible range and has a broad emission spectrum centered at approximately 690 nm (Sigma Aldrich, Saint Louis, MA, USA). As such, it can be used to simulate non-variable fluorescence emission from any source, including CDOM and free chlorophyll a. The BB3 was dissolved in MilliQ water to a final concentration of 118 µM. The volume of the BB3 spike was never more than 30 µL within each 20 mL sample. After spiking with BB3, each sample was dark-adapted for five minutes followed by a second test. In all cases, the F_b_ generated by addition of BB3 was at least three times the value of the original F_v_ and, consequently, decreased F_v_/F_m_ by approximately 65%.

### 2.11 Terminology

A structured approach has been taken in derivation of the parameters used within this manuscript. As baseline fluorescence is central to this study, new fluorescence terms to describe baseline-corrected values of existing fluorescence terms have been introduced. Otherwise, the parameters are structured around root terms that are widely used within the fluorescence community.

Table T1 provides terms used to describe the fluorescence signal at any point. Table T2 provides commonly used parameters derived from the terms in Table T1. Tables T3, T4 and T5 show the derivation of terms used for the yields, rate constants, absorption cross sections and absorption coefficients applied to PSII energy conversion processes. The remaining terms used are covered within Table T6.

The root terms and subscripts provided in Tables T3 and T4, respectively, are very widely used (examples include Butler and Kitajima, 1975; Kolber et al. 1998; Baker and Oxborough, 2004; and Oxborough et al. 2012). These tables were constructed to introduce consistency and minimize ambiguity: particularly with the distinction between absorption cross-sections and absorption coefficients. It should also be noted that the ‘optical absorption cross-section of PSII’ and ‘effective absorption cross-section of PSII’ (both unit area per photon) employed by Kolber et al. (1998) are, in terms of usage, equivalent to the absorption cross-sections of PSII light harvesting and PSII photochemistry (both unit area per PSII), respectively.

## 3 Results

### 3.1 Sample-specific K_a_ values under nutrient-replete and N-limited conditions

The sample-specific values of K_a_ for all nutrient-replete, low light grown cultures ranged from 7,822 m^−1^ for *C. vulgaris* to 25,743 m^−1^ for *T. pseudonana* (Figure 2A). Of the six species grown under both low and high light, only *D. salina* exhibited a significant difference in the K_a_ values between light treatments (Supplementary Table 1). In all cases, the N-limited sample-specific K_a_ values are significantly lower than for the nutrient-replete samples they were sub-cultured from (Figure 2A). These lower K_a_ values were matched to lower values of F_v_/F_m_ (Supplementary Table 1).

As previously discussed, there are two mechanisms that could cause sub-maximal F_v_/F_m_ values: dark-persistent Stern-Volmer quenching and baseline fluorescence. Importantly, the absorption method is insensitive to Stern-Volmer quenching while baseline fluorescence can introduce a significant error in the calculation of JV_PII_. In the context of these tests, the lower values of both sample-specific K_a_ and F_v_/F_m_ values observed within the N-limited cultures, when compared to the nutrient-replete values, are entirely consistent with a baseline fluorescence-induced error being introduced by, for example, the accumulation of photoinactivated PSII complexes. To test this possibility, Equation 14 (Oxborough, 2012) was used to derive a theoretical F_v_/F_mc_ value that could be applied across all N-limited cultures.

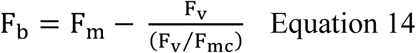

Where F_mc_ is the F_b_-corrected value of the measured F_m_ (see Terminology). When using this equation, F_m_ and F_v_ are measured from the sample and F_v_/F_mc_ is an assumed baseline corrected value of F_v_/F_m_ for the photochemically active PSII complexes within the sample (see Figure 1). The single, consensus value of F_v_/F_mc_ used was generated iteratively, by minimizing the total sum of squares for the differences in sample-specific K_a_ values from nutrient-replete cultures and corrected N-limited cultures.

**Figure 1.**
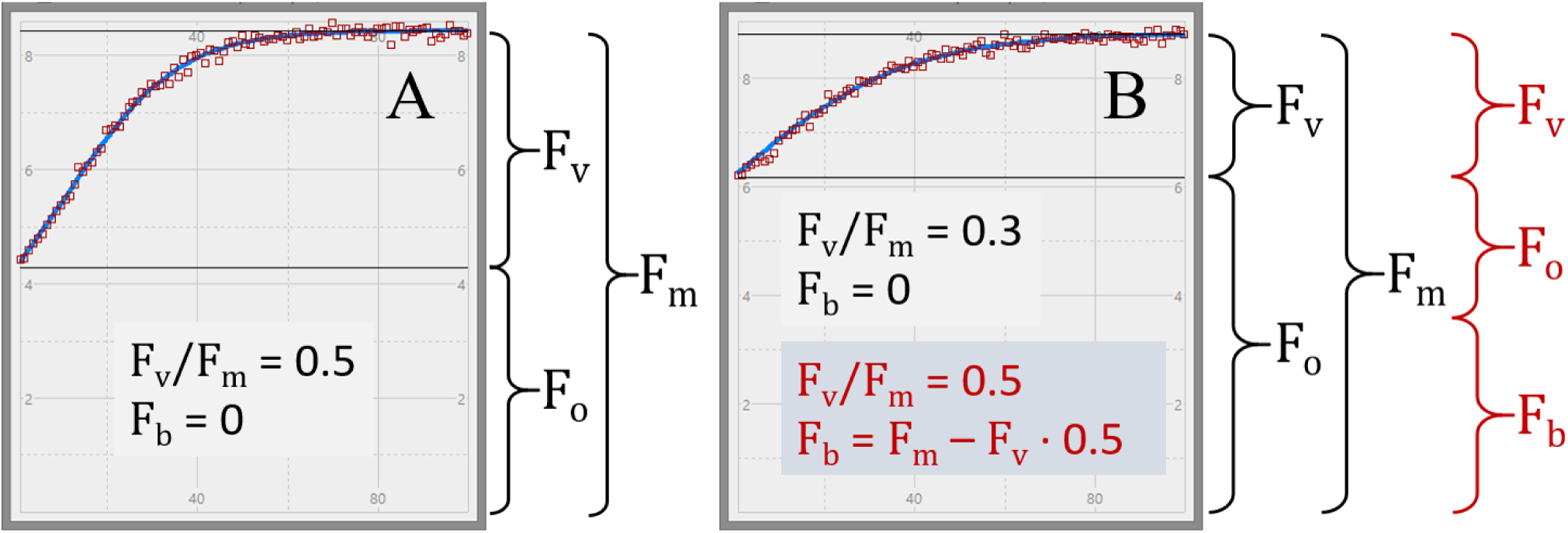
STAF measurements from *E. huxleyi* illustrating the concept of baseline fluorescence (F_b_). (A) is from a nutrient-replete culture in log-growth phase. (B) is from a N-limited culture. Two plausible explanations for the lower F_v_/F_m_ in (B) are considered within the main text. The first assumes that F_b_ is zero, as indicated by the black text in (B) and the second assumes that F_b_ has a value that accounts for the entire difference in F_v_/F_m_ between (A) and (B). All terms used are described within Terminology.

Applying an F_b_-correction within Equation 14 brings the N-limited sample-specific K_a_ values for all but one species (*T. pseudonana*) up to the point where they are not significantly different from the matched nutrient-replete culture values and resulted in a consensus F_v_/F_mc_ value of 0.518. Even with *T. pseudonana*, the F_b_-corrected value is much closer to the nutrient-replete value than is the uncorrected N-limited value. The observation that the consensus F_v_/F_mc_ value required for the F_b_-corrected values is slightly lower than most of the F_v_/F_m_ values measured from the nutrient-replete cultures (see Supplementary Table 1) may indicate that the photochemically active PSII complexes within the N-limited cultures are operating at a slightly lower efficiency that the photochemically active PSII complexes within the nutrient-replete cultures.

The dashed line within Figure 2A shows the default K_a_ value of 11,800 m^−1^ that is currently provided for the FastOcean sensor (hereafter, K_a_^FO^). Although this value falls within the mid-range of K_a_ values for the nutrient-replete cultures, there is considerable variability around this default value; for example, K_a_^FO^ is approximately 50% higher than the nutrient-replete, sample-specific K^a^ value for *C. vulgaris* and less than 50% of the equivalent value for *T. pseudonana*.

**Figure 2.**
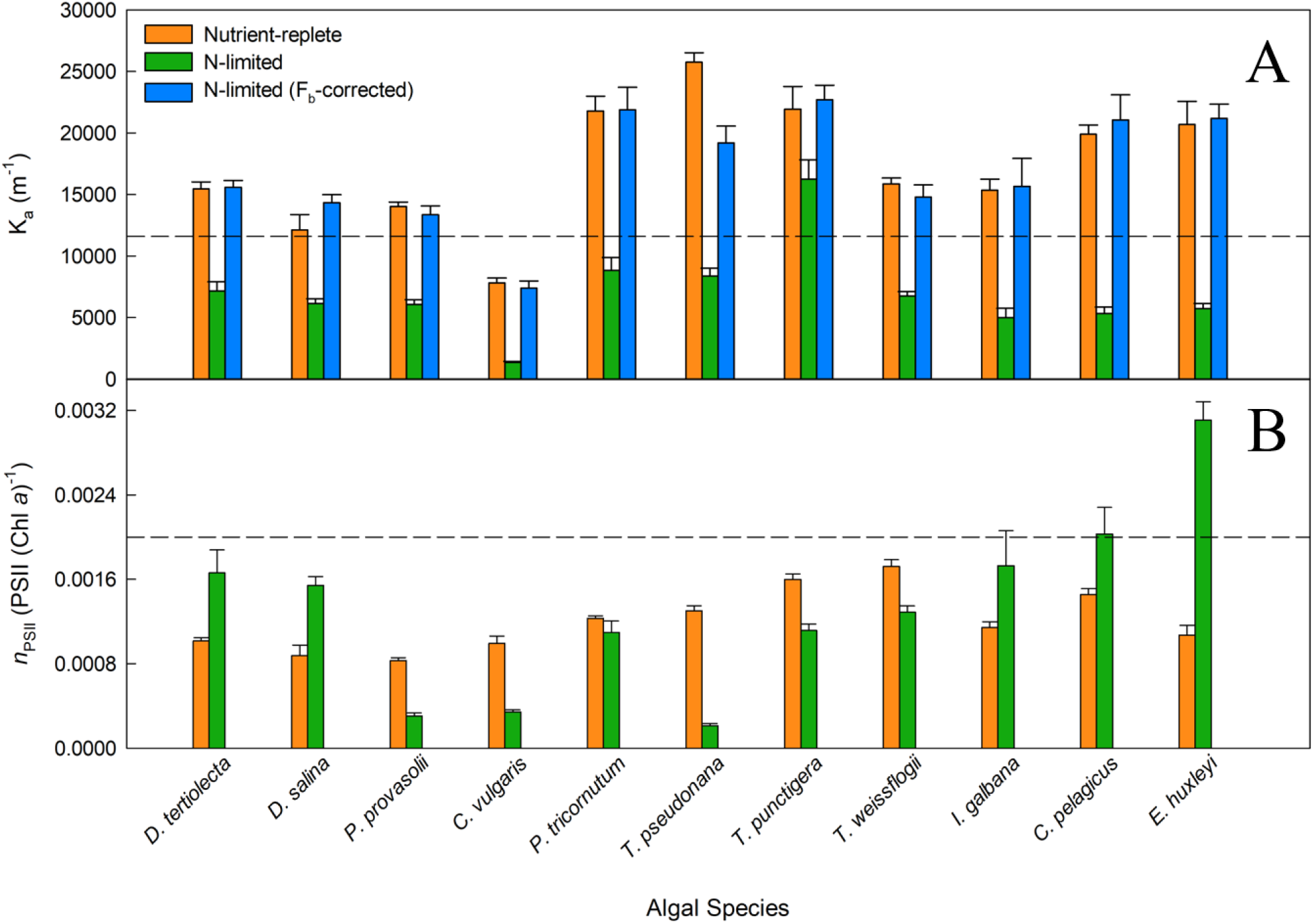
Variability in K_a_ (A) and *n*_PSII_ (B) values measured across a range of phytoplankton species. In (A), the consensus F_v_/F_mc_ value of 0.518 (see main text) was used to calculated F_b_ for each culture when the measured F_v_/F_m_ was lower than 0.518. F_b_ was subtracted from the measured F_o_ and F_m_. The dashed line represents K_a_^FO^ (11,800 m^−1^). Significant differences were tested by a series of parametric t-tests (t-test; P < 0.05); however, if normality was not achieved after data transformation a Mann-Whitney Rank Sum test was performed. All nutrient-replete and uncorrected N-limited K_a_ values were significantly different. In (B), the number of PSII reaction centers per chlorophyll a molecule was calculated from flash O_2_ measurements and chlorophyll *a* extractions. More details are provided within Materials and methods.

### 3.2 Interspecific variability in K_a_ and Chl PSII^−1^

A comparison between *n*_PSII_ and K_a_ is valid because they have a similar proportional impact in the calculation of [PSII] within Equations 2 and 4, respectively. Part B of Figure 2 shows *n*_PSII_ values for nutrient-replete cultures and N-limited cultures. The dashed line is at a widely used default value for *n*_PSII_ of 0.002 Chl PSII-1 (Kolber & Falkowski 1993; Suggett et al. 2001). There are two noteworthy features within this dataset. Firstly, the range of *n*_PSII_ values is very wide, at around 15:1: from less than 0.0002 Chl PSII^−1^ for the N-limited T. pseudonana to more than 0.003 Chl PSII^−1^ for N-limited *E. huxleyi*. Secondly, there is a lack of consistency between nPSII values from nutrient-replete cultures and N-limited cultures: five species show higher *n*_PSII_ values for the N-limited cultures while the remaining six species show lower *n*_PSII_ values for the N-limited cultures. Overall, these data provide a good illustration of how an assumed value for *n*_PSII_ can introduce large errors in the calculation of JV_PII_, which can only be corrected through independent determination of PSII concentration.

### 3.3 Comparison of OLC and FOC curves

Figure 3 shows OLC and FLC data from all eleven phytoplankton species used within this study. All data are from the nutrient-replete, low light-grown cultures. The FLC values of PhytoGO (y-axes) assume four electrons per O_2_ released. Values from the STAF data were derived using either K_a_^FO^ or the sample-specific values shown in Figure 2A. Clearly, in most cases, the match between OLC and FLC is greatly improved by using the sample-specific values of K_a_ in place of K_a_^FO^. The one exception is Figure 3B (*D. salina*) where the sample-specific K_a_ value happens to be very close to K_a_^FO^.

**Figure 3.**
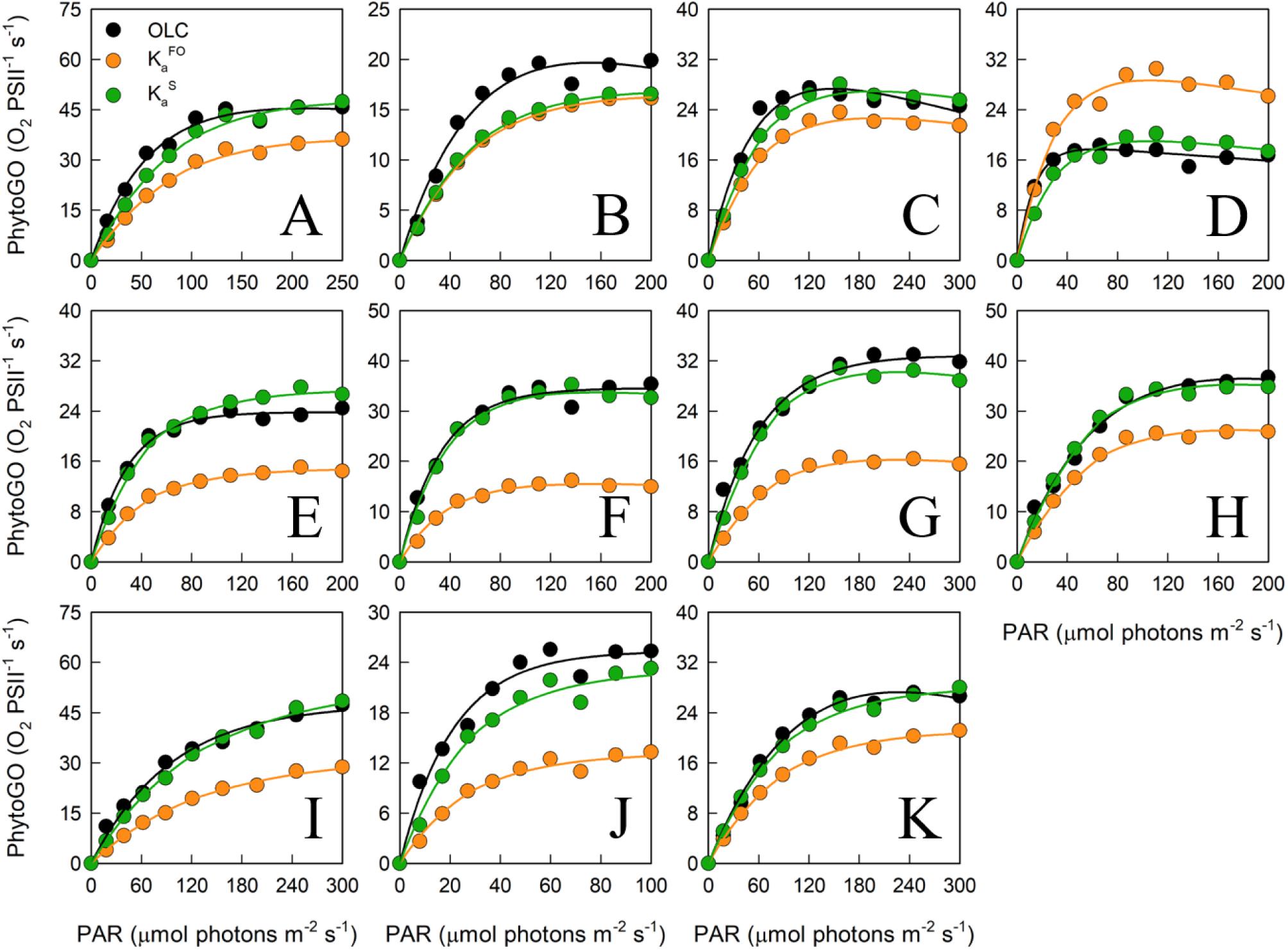
A representative example of simultaneous oxygen light curve (OLC) and fluorescence light curve (FLC) measurements made across all phytoplankton species. The OLC and FLC measurements were made on cultures acclimated to ambient temperature (∼ 20 °C) and low-light (LL = 30 µmol photons m^−2^ s^−1^). FLC data were standardized to equivalent units of O_2_, with both OLC and FLC data normalized to a derived concentration of functional PSII complexes (O_2_ PSII^−1^ s^−1^). FLC data were derived using K_a_^FO^ (11,800 m^−1^) or a sample-specific (K_a_^S^) value. The solid lines represent the P-E curve fits (color-coded to match the data points). Panel (A) = *D. tertiolecta*; (B) = *D. salina*; (C) = *P. provasolii*; (D) = *C. vulgaris*; (E) = *P. tricornutum*; (F) = *T. pseudonana*; (G) = *T. punctigera*; (H) = *T. weissflogii*; (I) = *C. pelagicus*; (J) = *E. huxleyi*; (K) = *I. galbana*. Replicates from each species (*n* = 5) are presented in Supplementary Figures 3 to 13.

The data presented within Figure 4 have been extracted from the OLCs and FLCs within Figure 3 to allow bulk comparison of the measured OLC and FLC PhytoGO values (A and C). Also shown is a comparison of the P_m_ values (B and D) from the OLC and FLC curve fits. Values were generated using either K_a_^FO^ (A and C) or the sample-specific values of K_a_ (B and D).

**Figure 4.**
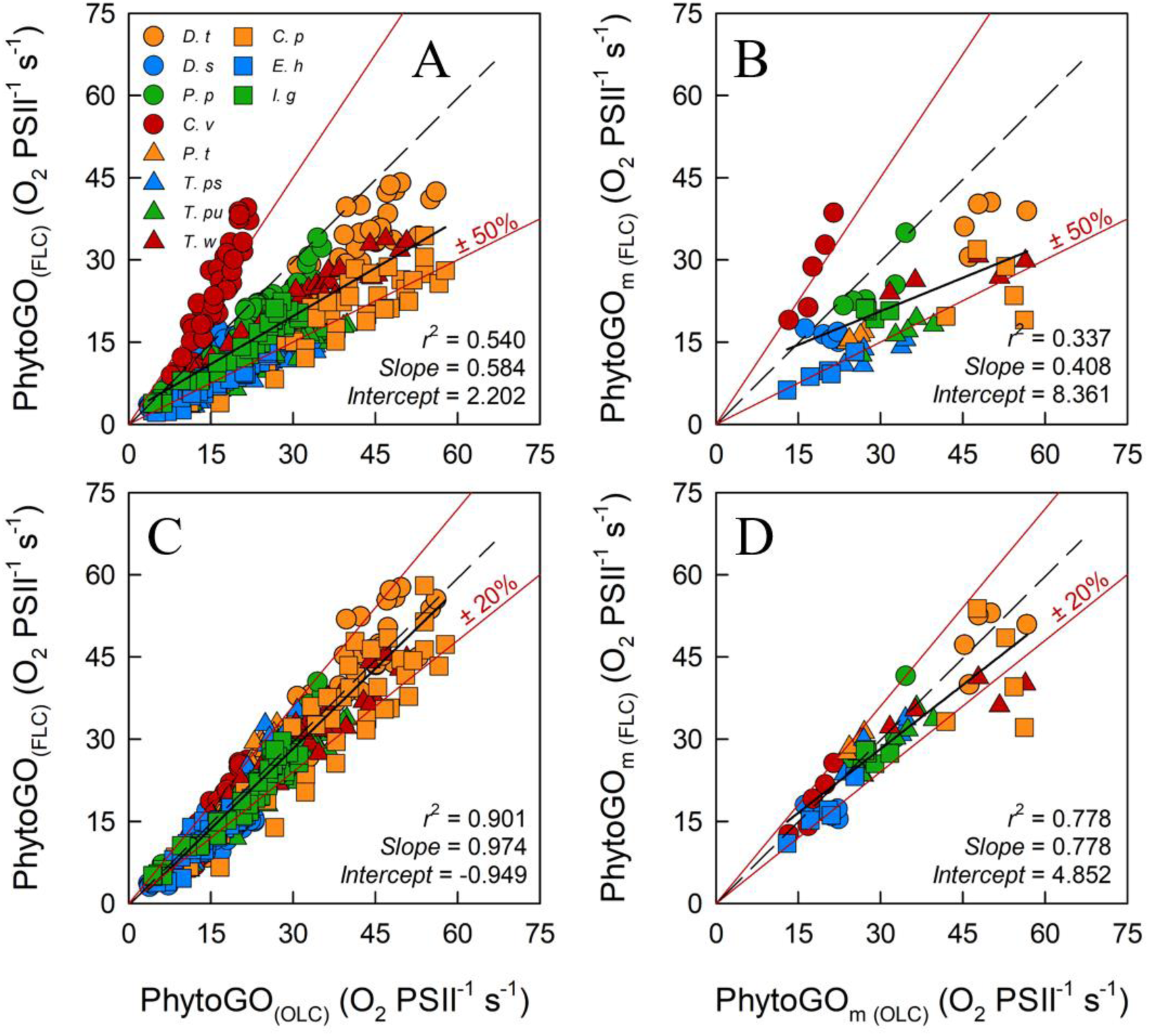
The relationship between the entire photosynthesis-photon irradiance (P-E) curve of PhytoGO, (A) and (C), and the maximum PhytoGO (PhytoGO_m_) from simultaneous OLC and FLC measurements, (B) and (D). FLC data were standardized to equivalent units of O_2_, with both OLC and FLC data normalized to a derived concentration of functional PSII complexes (O_2_ PSII^−1^ s^−1^). Within (A) and (B), FLC data were derived using K_a_^FO^ (11,800 m^−1^). Within (B) and (D), sample-specific values of K_a_ were used (see Materials and methods). Each species consisted of 5 biological replicates. The dashed line represents a 1:1 line, while the solid line is the linear regression used to generate r^2^, slope and intercept values. A key for the symbols in (A) is incorporated within Table 1.

Inevitably, the match between OLC and FLC data is improved significantly when the sample-specific values of K_a_ (C and D) are used in preference to K_a_^FO^ (A and B). The slope for the sample-specific K_a_ data is very close to the ‘ideal’ of 1.0 and a high proportion of data points fall within the ±20% lines included within the plot. In contrast, the K_a_^FO^ data have a much lower slope of 0.6 and ±50% lines are required to constrain a similar proportion of points. The sample-specific K_a_ values also generate a much better correlation between the values of P_m_ derived from OLC and FLC curve fits than K_a_^FO^ (D and B, respectively). The slope for the sample-specific K_a_ data (D), at 0.778, is significantly lower than the ideal of 1.0. This lower slope may be at least partly due to differences in the curve fits applied to OLC and FLC data (see Methods).

### 3.4 The stability of F_b_ under actinic light

Clearly, the consensus F_v_/F_mc_ (0.518) in Equation 14 generated a good match between K_a_ values for all but one of the nutrient-replete and N-limited cultures in Figure 2A. In a wider context, it could prove valid to use the consensus F_v_/F_mc_ of 0.518 within Equation 14 when the measured F_v_/F_m_ is lower than this value and assume that F_b_ is zero when the measured F_v_/F_m_ is above 0.518.

In situations where F_b_ is non-zero, the calculated value of a_LHII_ used within Equation 5 is decreased while value of F_q_’/F_m_’ used within the same equation is increased. The adjustment to a_LHII_ can largely be justified by the fact that matched F_b_ and a_LHII_ values are derived from the same dark-adapted STAF measurement. In contrast, the adjustment to F_q_’/F_m_’ is potentially more complex, simply because light-dependent NPQ can significantly decrease the maximum fluorescence level between the dark-adapted F_m_ and light-adapted F_m_’ (see Introduction). Given that a proportion of F_b_ could be from photoinactivated PSII complexes within the same membranes as the photochemically active PSII complexes, it seems reasonable to consider the possibility that NPQ could also quench F_b_.

To test the potential for a NPQ-dependent decrease in F_b_, additional FLCs were run on the N-limited cultures of *T. weissflogii* that had been sub-cultured from the low light-grown, nutrient-replete cultures. The value of F_b_ for the original, nutrient-replete cultures was always assumed to be zero, simply because the measured F_v_/F_m_ from was always above 0.518. Conversely, the F_v_/F_m_ values measured from the N-limited cultures were always well below the consensus value, at 0.116 ± 0.006.

Figure 5A shows the maximum PhytoGO values (PhytoGO_m_), measured as O_2_-evolution (x-axis) or calculated using the sample-specific K_a_ value from the nutrient-replete *T. weissflogii* of 15,868 m^−1^. For these values, F_b_ was set to zero for both the nutrient-replete cultures and the N-limited cultures. Clearly, while there is good agreement between the measured and calculated values of PhytoGO_m_ from the nutrient-replete cultures, most of the calculated PhytoGO_m_ values from the N-limited cultures are much higher than the measured values.

**Figure 5.**
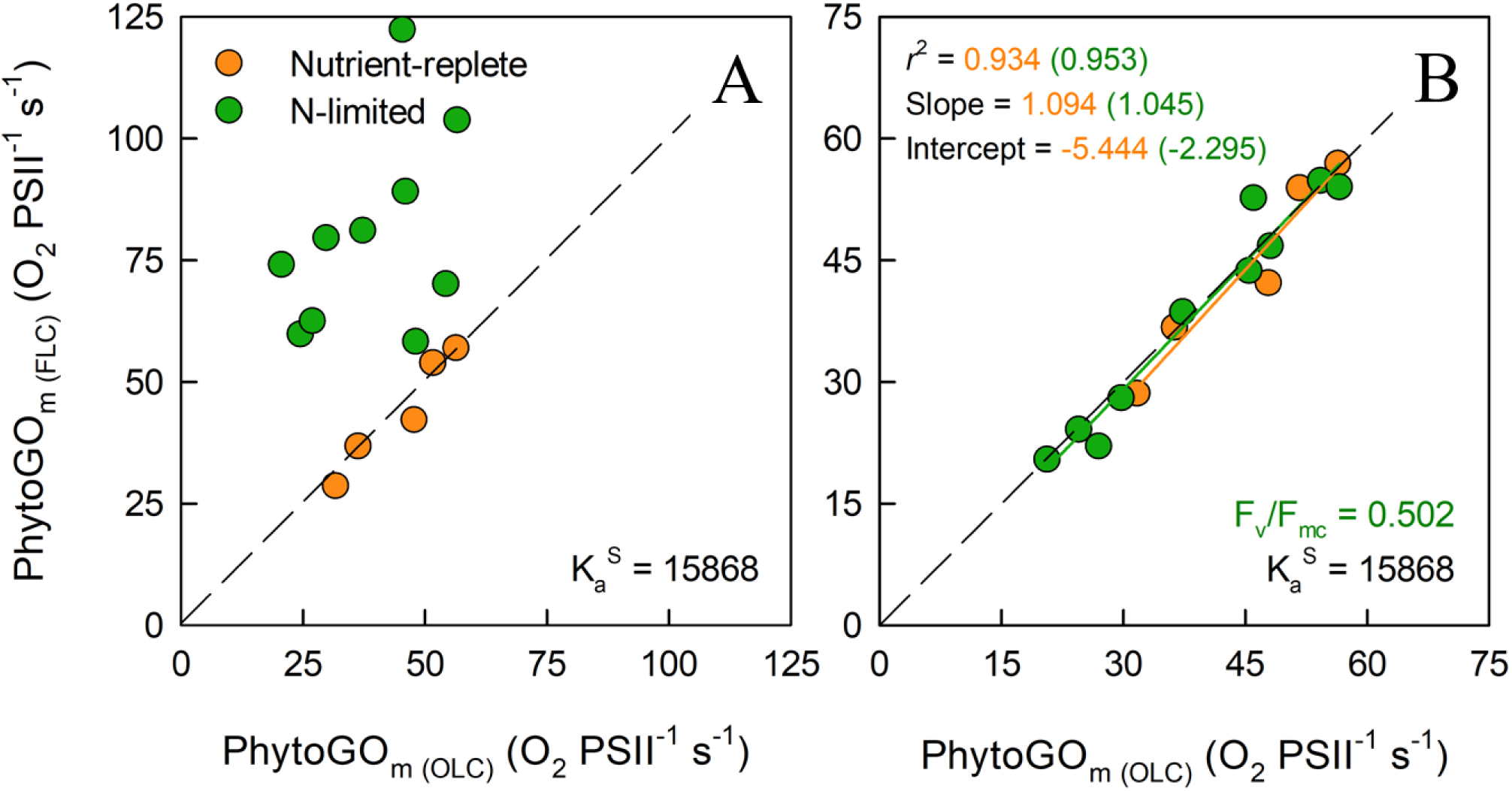
The relationship between simultaneous OLC and FLC measurements of maximum PhytoGO (PhytoGO_m_) within N-limited (*n* = 10) and nutrient replete (*n* = 5) *T. weissflogii* cultures. The sample-specific K_a_ value from the nutrient-replete cultures was applied throughout. In (A), no F_b_ correction was applied. In (B), the N-limited values were F_b_-corrected by applying a consensus F_v_/F_mc_ value of 0.502. Further details are provided within Materials and methods.

For Figure 5B, Equation 14 was used to generate a consensus F_v_/F_mc_ specific to the N-limited cultures. This consensus value was reached by minimizing the sum-of-squares for the regression line through the N-limited data by allowing F_b_ to vary. The mean consensus F_v_/F_mc_ from this fit (0.502) is within 3% of the consensus value derived from the dark-adapted data presented in Figure 2. In contrast, the average NPQ-dependent decrease from dark-adapted F_m_ to the light-adapted F_m_’ measured at P_m_ was always more than 30% (data not shown). Consequently, these data do not imply significant quenching of F_b_ between the dark-adapted state and P_m_.

### 3.5 Dual waveband STAF measurements to correct for the package effect

We hypothesized that the variance of sample-specific K_a_ values within Figure 2A could be at least partly due to variable package effect. As previously noted within Materials and methods, three FastBallast units (B730, B680 and B682) were used to measure fluorescence centered at 730 nm and 680 nm (both 10 nm FWHM) and 682 nm (30 nm FWHM), respectively.

To test the viability of a STAF-based approach to quantifying the package effect, we generated ratios of the F_v_ measured by B730 as a proportion of the F_v_ measured by B680 (F_v_^730/680^) or B683 (F_v_^730/683^). Within Figure 6, these values are plotted against sample-specific values of K_R_ (Figure 6A and D, respectively). The F_v_ ratios from Figure 6A and D were used to generate F_v_-derived values of K_R_ (Figure 6B and E, respectively).

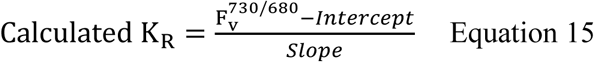

**Figure 6.**
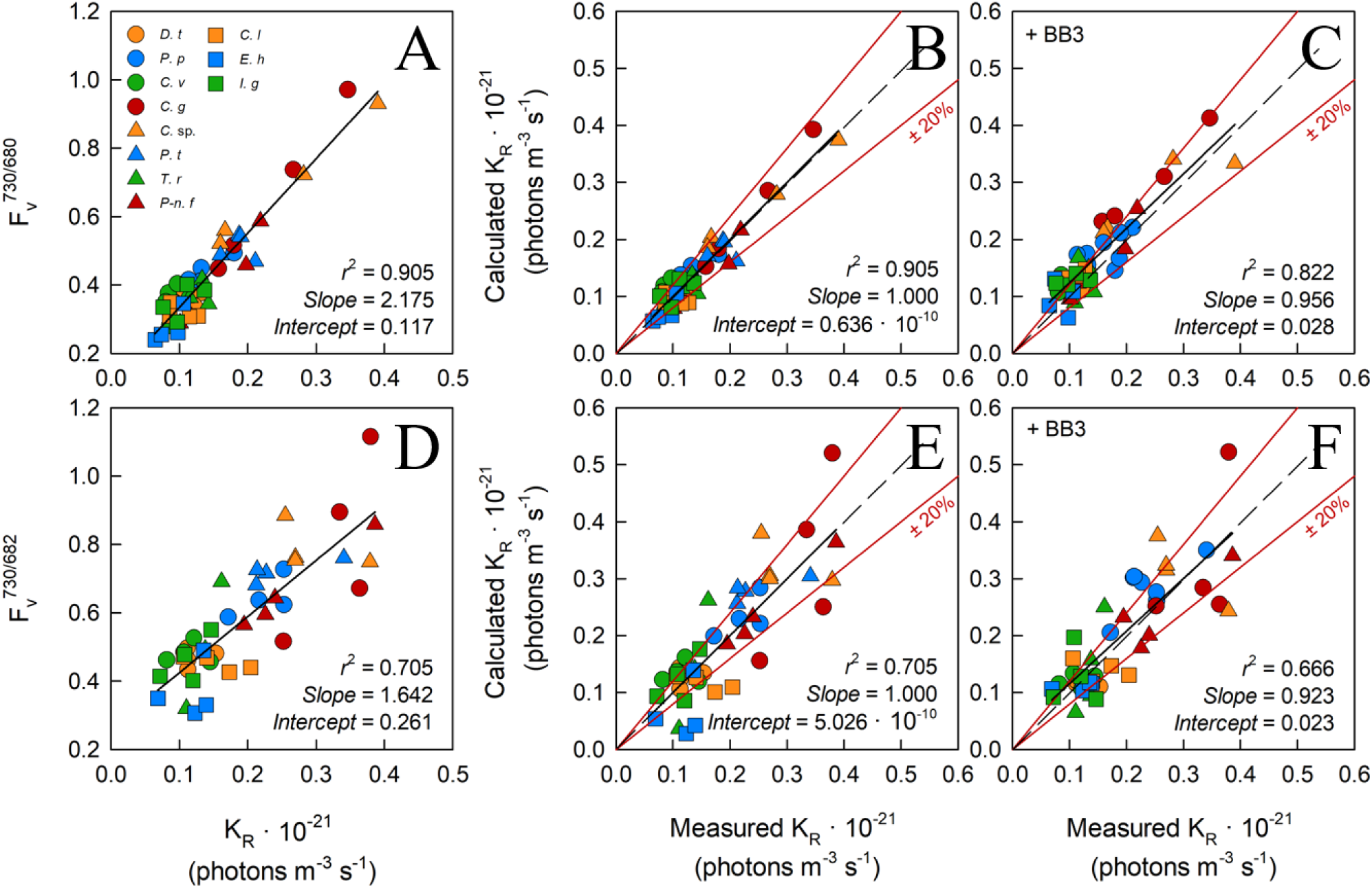
The relationship between dual wavelength STAF measurements and the sample-specific K_a_ values using the flash O_2_ method. The leftmost panels show the ratio of F_v_ measured by B730 to F_v_ measured by B680 (A) and F_v_ measured by B730 to F_v_ measured by B682 (D) against the measured K_a_ value from parallel measurements of flash O_2_. The middle panels, (B) and (E) show the relationship between flash O_2_-derived K_a_ and dual waveband-derived K_a_ values. The dual waveband K_a_ values within these panels were derived using the slopes and offsets reported in (A) and (D), as appropriate. The rightmost panels, (C) and (E), show the same relationships as reported in (B) and (E), respectively, but in the presence of extracellular baseline fluorescence (F_b_) generated by a spike of BB3 (see Materials and methods). The dashed lines in each panel represents a 1:1 slope, the solid black line is the linear regression. The solid red lines represent ± 20% of the regression values.

Equation 15 provides the conversion between A and D. For the equivalent conversion between B and E, F_v_^730/680^ was replaced with F_v_^730/683^. Slope and Intercept are the regression line values from A or D, as appropriate. The calculated K_R_ values within C and F were generated by combining the + BB3 F_v_^730/680^ and F_v_^730/683^ data with the Slope and Intercept from A and D, as appropriate.

One feature that is immediately clear from these data is the much tighter grouping of points along the regression lines for the F_v_^730/680^ data (A to C) than the F_v_^730/683^ data (D to F). This indicates that the 30 nm FWHM of the 682 nm bandpass filter is too broad to adequately isolate the fluorescence generated close the 680 nm absorption peak and, consequently, that the 10 nm FWHM 680 nm bandpass filter is the better choice for these measurements.

All 11 species used within the package effect tests were grown under nutrient-replete conditions and exhibited F_v_/F_m_ values that were above the consensus value of 0.518 generated from the first part of this study. The addition of BB3 to each sample within the package effect tests was to simulate the lower F_v_/F_m_ values that are frequently observed under conditions of stress. The expectation was that fluorescence from the added BB3 would increase F_b_ but have minimal impact F_v_ and, as a consequence, that the slope of the relationship between calculated and measured K_R_ values would not be significantly affected by a BB3-dependent increase in F_b_. The absence of significant changes in slope between B and C and E and F are consistent with this expectation.

### 3.6 Discussion

The absorption method described by Oxborough et al. (2012) provides a method for estimating PhytoGO and PhytoPP on much wider spatiotemporal scales than O_2_ evolution or ^14^C fixation, respectively, through determination of JV_PII_. This study was undertaken to assess the extent to which baseline fluorescence and the package effect could introduce errors into the calculation of JV_PII_ (Equation 5).

With regard to baseline fluorescence (F_b_), the underlying question was whether sub-maximal dark-adapted value of F_v_/F_m_ could be attributed to F_b_ or downregulation of PSII photochemistry by dark-persistent Stern-Volmer quenching or some combination of the two. The data presented within Figure 2A provides strong evidence that, for the examples presented within this study, F_b_ is by far the dominant contributor to sub-maximal F_v_/F_m_ values. Although this interpretation may not hold for all phytoplankton species and environmental conditions, this study provides a straightforward, practical approach to addressing the question of how universally valid an F_b_ correction to low sub-maximal dark-adapted F_v_/F_m_ values might be.

We conclude that no correction for baseline fluorescence should be applied when the dark-adapted F_v_/F_m_ is above a certain consensus value. In situations where the dark-adapted F_v_/F_m_ is below this consensus value, Equation 14 should be used to calculate a value for F_b_. From the data presented here, a consensus value (F_v_/F_mc_) of between 0.50 and 0.52 seems an appropriate default value.

Clearly, the value of F_b_ generated by Equation 14 is dependent on a STAF measurement made on a dark-adapted sample. The data presented in Figure 5 indicate that, for this specific example at least, the dark-adapted F_b_ could be applied at the other end of the FLC scale, to correct the value of P_m_.

With regard to the package effect, the wide range of K_a_ values within Figure 2A is entirely consistent with a significant proportion of the fluorescence emitted from functional PSII complexes being reabsorbed through this process. This interpretation is clearly supported by data presented in Figure 3, where use of the sample-specific K_a_ value in place of K_a_^FO^ provides a much stronger match between the FLC and OLC data. The dual waveband data presented in Figure 6 provide strong evidence that the package effect-induced error could be decreased significantly through incorporation of a F_v_^730/680^-derived correction factor applied to a default instrument-type specific K_a_ value such as K_a_^FO^. From a practical point of view, routine implementation of this correction step will require either two detectors with different filters or a single detector with switchable filtering. On balance, the latter option is likely to prove more cost-effective and easier to calibrate.

Overall, the conclusions reached can be summarized by Equation 16

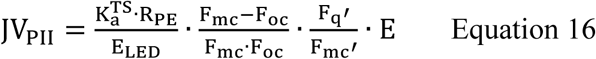

where K_a_^TS^ is the instrument type-specific K_a_ value and R_PE_ is a dimensionless sample-specific correction factor. All other terms are as before or are defined in Terminology.

For the species and conditions examined in this paper, the data presented provide strong evidence that baseline correction and package effect correction can increase the accuracy of estimates of PhytoGO from STAF. We anticipate that development and deployment of STAF instrumentation that will allow Equation 16 to be applied will take us significantly closer to achieving the objective of obtaining reliable autonomous estimates of PhytoGO. Such measurements, if used in conjunction with simultaneous satellite measurements of ocean color, will likely lead to improved estimates of local, regional or global pelagic PhytoPP.

## 3.7 Acknowledgements

The authors wish to thank Tania Cresswell-Maynard, James Fox and Philipp Siegel (University of Essex, UK) for supplying the phytoplankton cultures. We would also like to thank Hoi Ga Chan (University of Princeton, US) and Mark Moore and Anna Hickman (Southampton University, UK) for helpful discussions. RJG acknowledges support from NERC grant NE/P002374/1.

## 3.8 Author contributions statement

KO conceived of the study and developed the software required to conduct the experiments. The initial baseline experiments were designed by KO, RG and TB. All of the baseline experiments were conducted by TB who also processed the primary data. The dual waveband experiments for assessment of the package effect were conceived of by KO and conducted by TB. Package effect data were processed by TB and jointly analyzed by KO and TB. Figure 1 was produced by KO, all remaining Figures were produced by TB. The initial draft of the main text was produced by KO. Iterations of the manuscript were implemented by KO, TB and RG. The submitted version of the manuscript is approved for publication by KO, TB and RG.

## 3.10 Contribution to the field statement

Phytoplankton photosynthesis is responsible for approximately half of the carbon fixed on the planet. As a process, photosynthesis is responsive to variability in multiple environmental drivers including light, temperature and nutrients across spatial scales from meters to ocean basins, and time scales from minutes to tens of years. This poses significant challenges for measurement and monitoring. While direct measurement of the carbon fixed by photosynthesis can only be applied on very limited spatial and temporal scales, active chlorophyll fluorescence has enormous potential for the accurate measurement of phytoplankton photochemistry, which provides the reducing power for carbon fixation, on much wider spatiotemporal scales and with much lower operational costs. This study identifies practical measures that can be taken to improve the accuracy of such measurements. We are confident that these measures will have minimal impact on the frequency at which phytoplankton photochemistry is assessed and that they will be suitable for application on autonomous measurement platforms.

---

**Table T1.** The o, m and v subscripts define the origin (of variable fluorescence), maximum fluorescence and variable fluorescence, respectively. The q subscript defines the proportion of variable fluorescence that is quenched by PSII photochemistry. The b subscript defines baseline fluorescence, which is assumed to contribute equally to F_o_, F_m_ and F. In the interest of readability, only dark-adapted values have been included in the Measurement or derivation column.

| Dark term | Light term | Measurement or derivation |
| --- | --- | --- |
| $F_o$ | $F_o'$ | Extrapolation to $t = 0$ from a ST pulse |
| $F_m$ | $F_m'$ | At the asymptote of a ST pulse |
| $F_v$ | $F_v'$ | $F_m - F_o$ |
| $F$ | $F'$ | Any point between $F_o$ and $F_m$ |
| $F_q$ | $F_q'$ | $F_m - F$ |
| $F_b$ | $F_b'$ | Fluorescence signal not attributable to functional PSII centers |
| $F_b$ | $F_b'$ | Proportion of the measured $F_o$ that does not arise from functional PSII centers in the open state |
| $F_{oc}$ | $F_{oc}'$ | The baseline subtracted value of $F_o$ such that $F_{oc} = F_o - F_b$ |
| $F_{mc}$ | $F_{mc}'$ | The baseline subtracted value of $F_m$ such that $F_{mc} = F_m - F_b$ |
| $F_c$ | $F_c'$ | The baseline subtracted value of $F$ such that $F_c = F - F_b$ |

**Table T2.** Fluorescence parameters derived using the terms within Table T1.

| Dark parameter | Light parameter | Interpretation |
| --- | --- | --- |
| $F_v/F_m$ | $F_v'/F_m'$ | Provides an estimate of PSII photochemical efficiency ($\phi_{PII}$) when all PSII centers are in the open state and $F_b = 0$ |
| $F_v/F_{mc}$ | $F_v'/F_{mc}'$ | Provides a baseline-corrected estimate of PSII photochemical efficiency ($\phi_{PII}$) when all PSII centers are in the open state |
| $F_q/F_m$ | $F_q'/F_m'$ | Provides an estimate of PSII photochemical efficiency ($\phi_{PII}$) when some centers are closed and $F_b = 0$ |
| $F_q/F_{mc}$ | $F_q'/F_{mc}'$ | Provides a baseline-corrected estimate of PSII photochemical efficiency ($\phi_{PII}$) when some centers are closed |
| $F_q/F_v$ | $F_q'/F_v'$ | Provides a value for the PSII photochemical factor |
| $F_o/F_v$ | $F_o'/F_v'$ | Provides an estimate of Stern-Volmer quenching within the PSII pigment matrix, normalized to PSII photochemistry (only valid when $F_b = 0$) |
| $F_{oc}/F_v$ | $F_{oc}'/F_v'$ | Provides a baseline-corrected estimate of Stern-Volmer quenching within the PSII pigment matrix, normalized to PSII photochemistry |

**Table T3.** Root terms used in the derivation of parameter ‘x’ within Table T5.

| Term | Meaning | Units |
| --- | --- | --- |
| $\phi_x$ | Yield | dimensionless |
| $k_x$ | Rate constant | photons s<sup>-1</sup> |
| $\sigma_x$ | Absorption cross-section | m<sup>-2</sup> PSII<sup>-1</sup> |
| $a_x$ | Absorption coefficient | m<sup>-1</sup> |

**Table T4.** Subscripts used for derivation of parameters within Table T5.

| Term | Usage |
| --- | --- |
| II | Photosystem II (PSII) |
| LH | Light-harvesting |
| P | Photochemistry |
| F | Fluorescence |
| D | Non-radiative decay |

**Table T5.** Parameters derived from the root terms and subscripts within Tables T3 and T4, respectively. Empty fields within the Light term column indicate an assumed lack of change for these quantities between the dark and light-adapted states.

| Dark term | Light term | Definition | SI units |
| --- | --- | --- | --- |
| $\phi_{PII}$ | $\phi_{PII}'$ | Yield of PSII photochemistry | dimensionless |
| $\phi_{FII}$ | $\phi_{FII}'$ | Yield of PSII fluorescence | dimensionless |
| $\phi_{DII}$ | $\phi_{DII}'$ | Yield of non-radiative decay within PSII | dimensionless |
| $k_{PII}$ | | Rate constant for photochemistry at PSII | photons s<sup>-1</sup> |
| $k_{FII}$ | | Rate constant for fluorescence emission from PSII | photons s<sup>-1</sup> |
| $k_{DII}$ | $k_{DII}'$ | Rate constant for non-radiative decay within PSII | photons s<sup>-1</sup> |
| $\sigma_{LHII}$ | | Absorption cross-section of PSII light harvesting | m<sup>2</sup> PSII<sup>-1</sup> |
| $\sigma_{PII}$ | $\sigma_{PII}'$ | Absorption cross-section of PSII photochemistry | m<sup>2</sup> PSII<sup>-1</sup> |
| $\sigma_{FII}$ | $\sigma_{FII}'$ | Absorption cross-section of PSII fluorescence | m<sup>2</sup> PSII<sup>-1</sup> |
| $\sigma_{DII}$ | $\sigma_{DII}'$ | Absorption cross-section of PSII non-radiative | m<sup>2</sup> PSII<sup>-1</sup> |
| $a_{LHII}$ | | Absorption coefficient of PSII light harvesting | m<sup>-1</sup> |
| $a_{PII}$ | $a_{PII}'$ | Absorption coefficient of PSII photochemistry | m<sup>-1</sup> |
| $a_{FII}$ | $a_{FII}'$ | Absorption coefficient of PSII fluorescence | m<sup>-1</sup> |
| $a_{DII}$ | $a_{DII}'$ | Absorption coefficient of PSII non-radiative decay | m<sup>-1</sup> |

