## Supplemental Tables and Figures for "Improving the accuracy of single turnover active fluorometry (STAF) for the estimation of phytoplankton primary productivity (PhytoPP)"

**Supplementary Table 1.** The cell dimensions, dark-adapted $F_v/F_m$ and sample-specific $K_a$ values of the 11 phytoplankton cultures used for the baseline fluorescence experiments.

| *Algal species* | Cell characteristics | | | $F_v/F_m$ | | | $K_a$ | | |
|---|---|---|---|---|---|---|---|---|---|
| | Width (µm) | Length (µm) | Volume ($\mu m^3$) | HL | LL | N-Limited | HL | LL | N-Limited |
| *C. vulgaris* | 2.4 (0.1) | 2.9 (0.1) | 9 (1) | - | 0.535 (0.003) | 0.156 (0.004) | - | 7822 (371) | 1346 (96) |
| *C. pelagicus* | 10.0 (0.3) | 10.0 (0.3) | 567 (46) | - | 0.538 (0.004) | 0.217 (0.004) | - | 19898 (678) | 5331 (528) |
| *D. salina* | 10.9 (0.1) | 13.4 (0.1) | 853 (26) | 0.543 (0.008) | 0.539 (0.008) | 0.313 (0.032) | 10735 (270) | 12117 (1477) | 6155 (368) |
| *D. tertiolecta* | 11.4 (0.4) | 16.3 (0.3) | 1119 (46) | 0.515 (0.009) | 0.528 (0.004) | 0.320 (0.042) | 16400 (736) | 15455 (598) | 7148 (774) |
| *E. huxleyi* | 4.0 (0.2) | 4.0 (0.2) | 36 (6) | 0.511 (0.002) | 0.533 (0.002) | 0.215 (0.009) | 21978 (1255) | 20677 (1819) | 5732 (413) |
| *I. galbana* | 3.5 (0.1) | 4.7 (0.1) | 33 (2) | - | 0.534 (0.003) | 0.255 (0.005) | - | 15352 (888) | 5001 (755) |
| *P. provasolii* | 5.8 (0.1) | 7.3 (0.1) | 139 (7) | - | 0.532 (0.003) | 0.318 (0.010) | - | 14032 (330) | 6075 (378) |
| *P. tricornutum* | 3.1 (0.1) | 38.3 (0.6) | 145 (10) | 0.519 (0.005) | 0.561 (0.007) | 0.298 (0.048) | 21456 (454) | 21796 (931) | 8823 (1051) |
| *T. pseudonana* | 7.3 (0.1) | 8.8 (0.1) | 234 (10) | - | 0.544 (0.006) | 0.304 (0.019) | - | 25743 (735) | 8372 (648) |
| *T. punctigera* | 30.6 (0.2) | 44.9 (0.2) | 68875 (5171) | 0.519 (0.002) | 0.545 (0.007) | 0.426 (0.050) | 25450 (909) | 21978 (1255) | 16248 (1575) |
| *T. weissflogii* | 17.5 (0.1) | 24.3 (0.1) | 4111 (55) | 0.521 (0.005) | 0.537 (0.003) | 0.333 (0.017) | 17228 (602) | 15868 (320) | 6744 (358) |

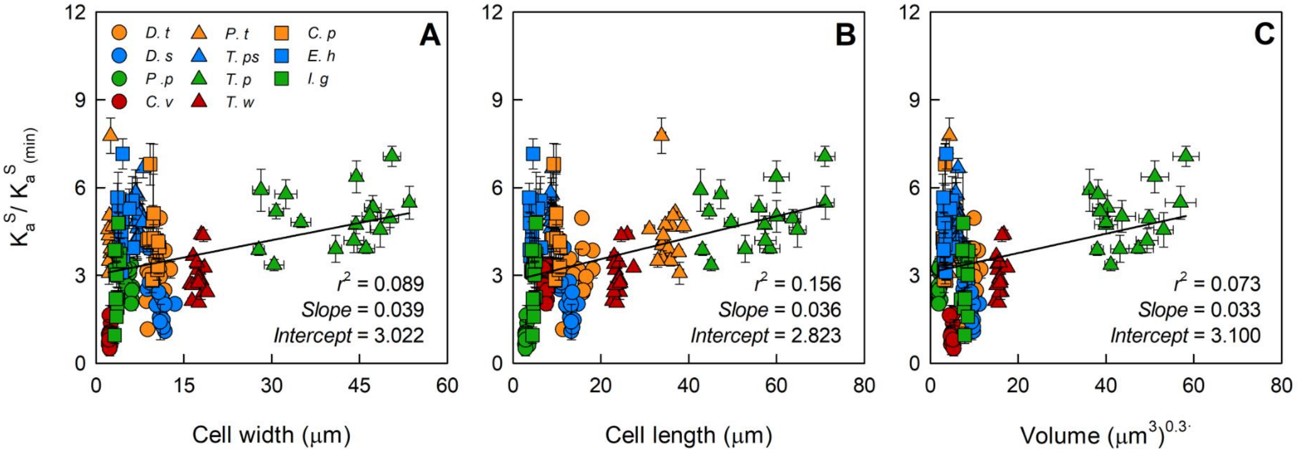


**Supplementary Figure 1.** The relationship between sample-specific $K_a$ values normalized to the lowest measured sample-specific $K_a$ value and cell width (A), cell length (B) or cell volume (C). The solid line is the linear regression used to generate the $r^2$, slope and intercept values. *D. t = D. tertiolecta; D. s = D. salina; P. p = P. provasolii; C. v = C. vulgaris; P. t = P. tricornutum; T. ps = T. pseudonana; T. pu = T. punctigera; T. w = T. weissflogii; C. p = C. pelagicus; E. h = E. huxleyi; I. g = I. galbana.*

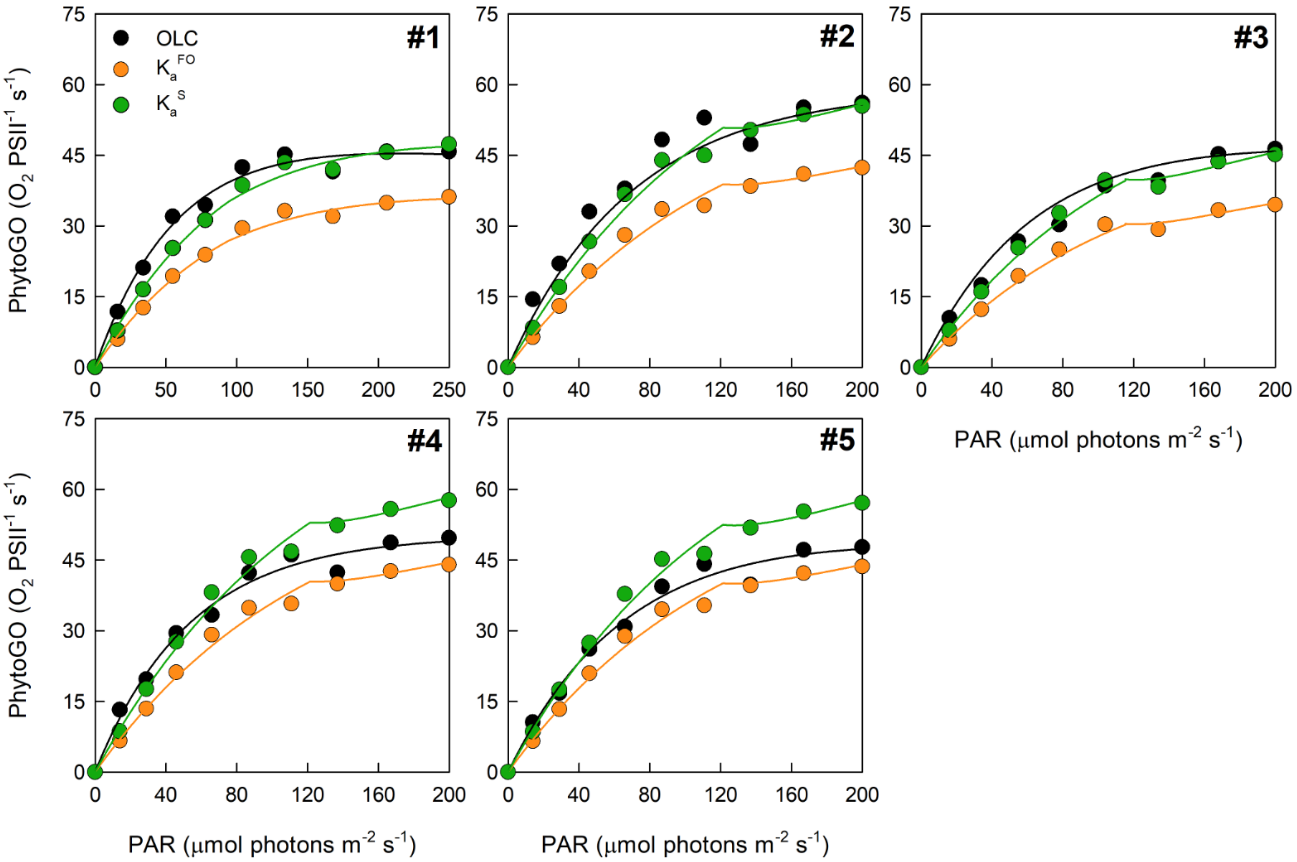


**Supplementary Figure 2.** The replicate simultaneous oxygen light curve (OLC) and fluorescence light curve (FLC) measurements made on *D. tertiolecta.* The OLC and FLC measurements were made on cultures acclimated to ambient temperature (~ 20 °C) and low-light (LL = 30 μmol photons $m^{-2}$ $s^{-1}$). FLC data was standardized to equivalent units of $O_2$, with both OLC and FLC data normalized to a derived concentration of PSII reaction centers (RCII) ($O_2$ RCII $s^{-1}$). FLC data were calculated using $K_a^{FO}$ = 11,800 $m^{-1}$ or the sample-specific $K_a^{S}$ value of 15,455 $m^{-1}$. The solid lines represent the PE curve fits. Each panel represents a biological replicate.

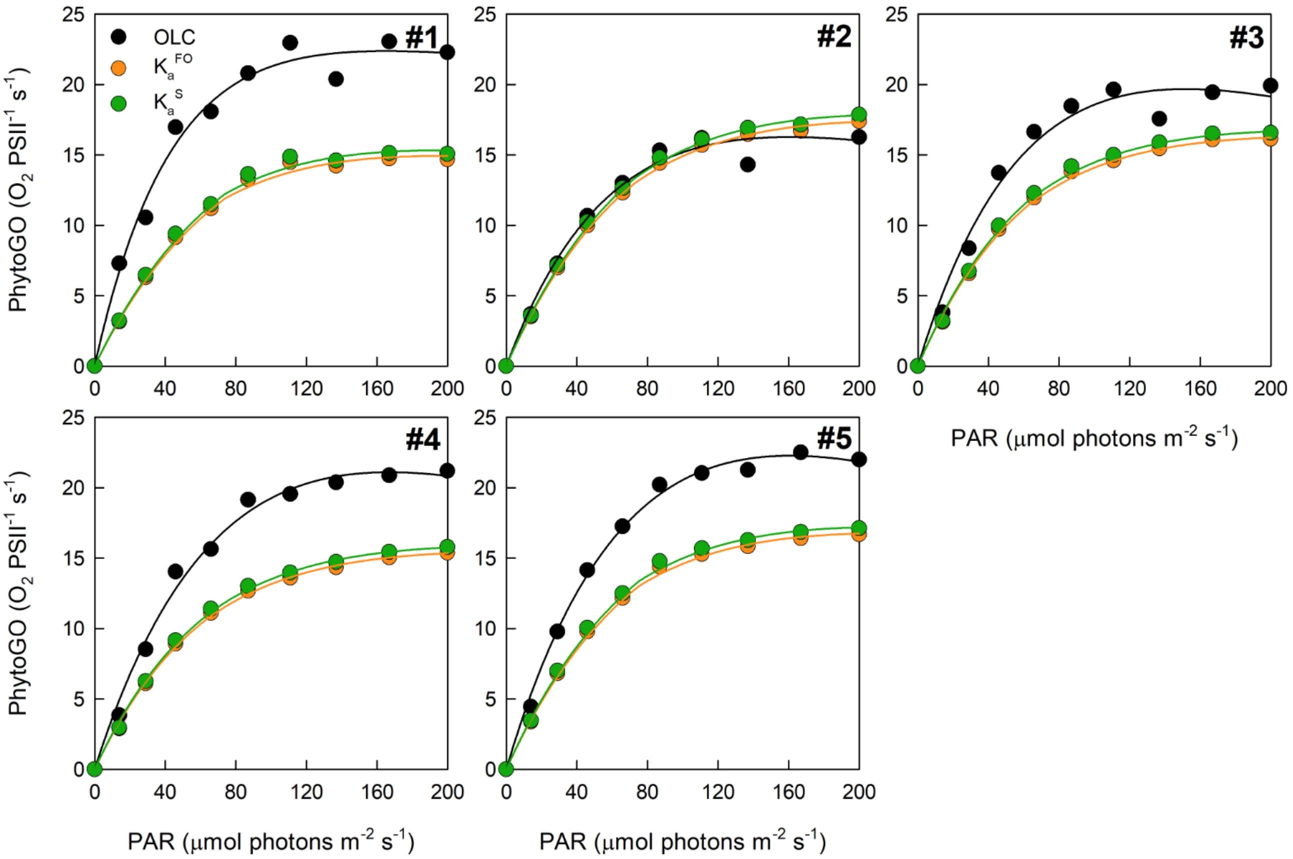


**Supplementary Figure 3.** The replicate simultaneous oxygen light curve (OLC) and fluorescence light curve (FLC) measurements made on *D. salina*. The OLC and FLC measurements were made on cultures acclimated to ambient temperature (~ 20 °C) and low-light (LL = 30 μmol photons $m^{-2}$ $s^{-1}$). FLC data was standardized to equivalent units of $O_2$, with both OLC and FLC data normalized to a derived concentration of PSII reaction centers (RCII) ($O_2$ RCII $s^{-1}$). FLC data were calculated using $K_a^{FO}$ = 11,800 $m^{-1}$ or the sample-specific $K_a^S$ value of 12,117 $m^{-1}$. The solid lines represent the PE curve fits. Each panel represents a biological replicate.

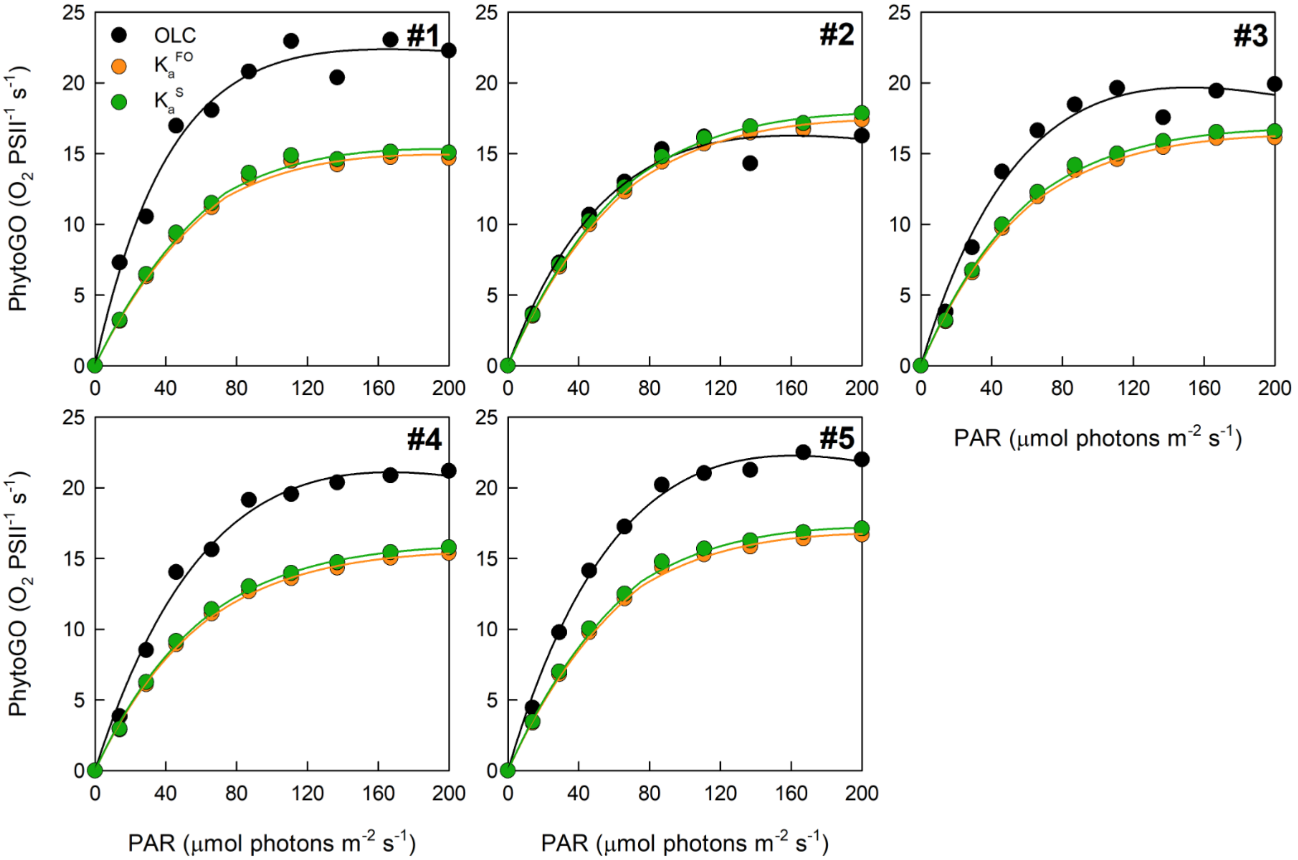


**Supplementary Figure 4.** The replicate simultaneous oxygen light curve (OLC) and fluorescence light curve (FLC) measurements made on *P. provasolii*. The OLC and FLC measurements were made on cultures acclimated to ambient temperature (~ 20 °C) and low-light (LL = 30 µmol photons m$^{-2}$ s$^{-1}$). FLC data was standardized to equivalent units of $O_2$, with both OLC and FLC data normalized to a derived concentration of PSII reaction centers (RCII) ($O_2$ RCII s$^{-1}$). FLC data were calculated using $K_a^{FO}$ = 11,800 m$^{-1}$ or the sample-specific $K_a^S$ value of 14,032 m$^{-1}$. The solid lines represent the PE curve fits. Each panel represents a biological replicate.

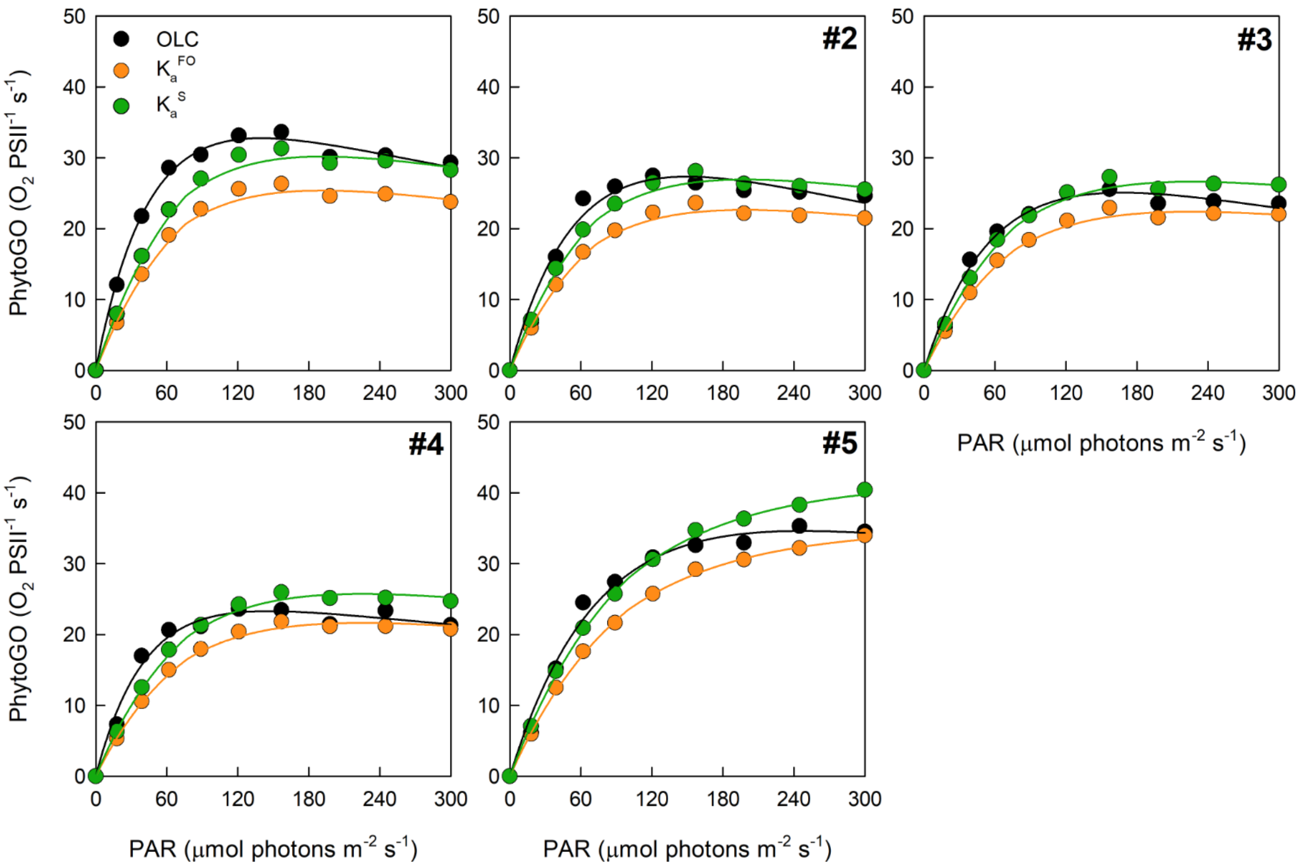


**Supplementary Figure 5.** The replicate simultaneous oxygen light curve (OLC) and fluorescence light curve (FLC) measurements made on *C. vulgaris*. The OLC and FLC measurements were made on cultures acclimated to ambient temperature (~ 20 °C) and low-light (LL = 30 μmol photons $m^{-2}$ $s^{-1}$). FLC data was standardized to equivalent units of $O_2$, with both OLC and FLC data normalized to a derived concentration of PSII reaction centers (RCII) ($O_2$ RCII $s^{-1}$). FLC data were calculated using $K_a^{FO}$ = 11,800 $m^{-1}$ or the sample-specific $K_a^S$ value of 7,822 $m^{-1}$. The solid lines represent the PE curve fits. Each panel represents a biological replicate.

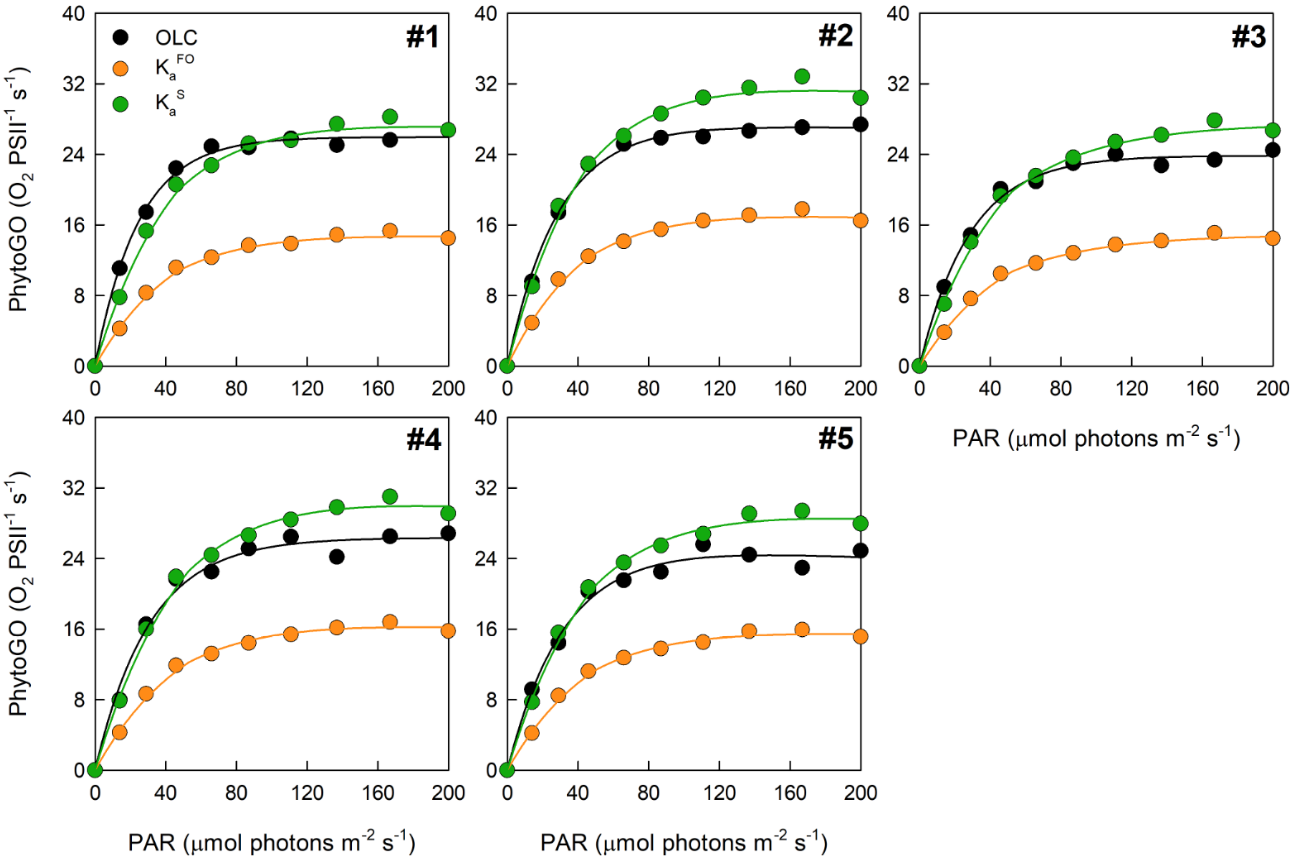


**Supplementary Figure 6.** The replicate simultaneous oxygen light curve (OLC) and fluorescence light curve (FLC) measurements made on *P. tricornutum*. The OLC and FLC measurements were made on cultures acclimated to ambient temperature (~ 20 °C) and low-light (LL = 30 μmol photons m$^{-2}$ s$^{-1}$). FLC data was standardized to equivalent units of $O_2$, with both OLC and FLC data normalized to a derived concentration of PSII reaction centers (RCII) ($O_2$ RCII s$^{-1}$). FLC data were calculated using $K_a^{FO}$ = 11,800 m$^{-1}$ or the sample-specific $K_a^{S}$ value of 21,796 m$^{-1}$. The solid lines represent the PE curve fits. Each panel represents a biological replicate.

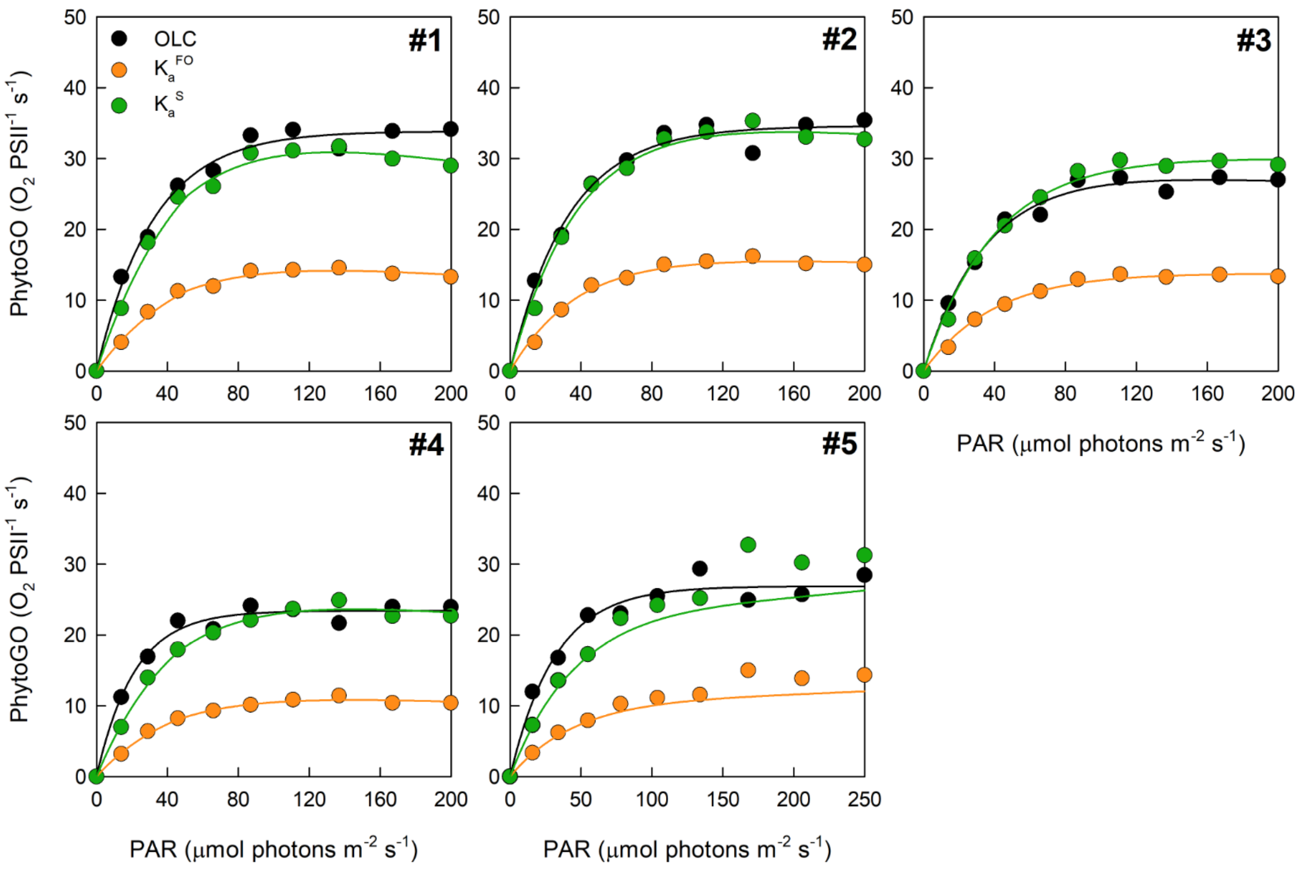


**Supplementary Figure 7.** The replicate simultaneous oxygen light curve (OLC) and fluorescence light curve (FLC) measurements made on *T. pseudonana*. The OLC and FLC measurements were made on cultures acclimated to ambient temperature (~ 20 °C) and low-light (LL = 30 µmol photons $m^{-2}$ $s^{-1}$). FLC data was standardized to equivalent units of $O_2$, with both OLC and FLC data normalized to a derived concentration of PSII reaction centers (RCII) ($O_2$ RCII $s^{-1}$). FLC data were calculated using $K_a^{FO}$ = 11,800 $m^{-1}$ or the sample-specific $K_a^S$ value of 25,743 $m^{-1}$. The solid lines represent the PE curve fits. Each panel represents a biological replicate.

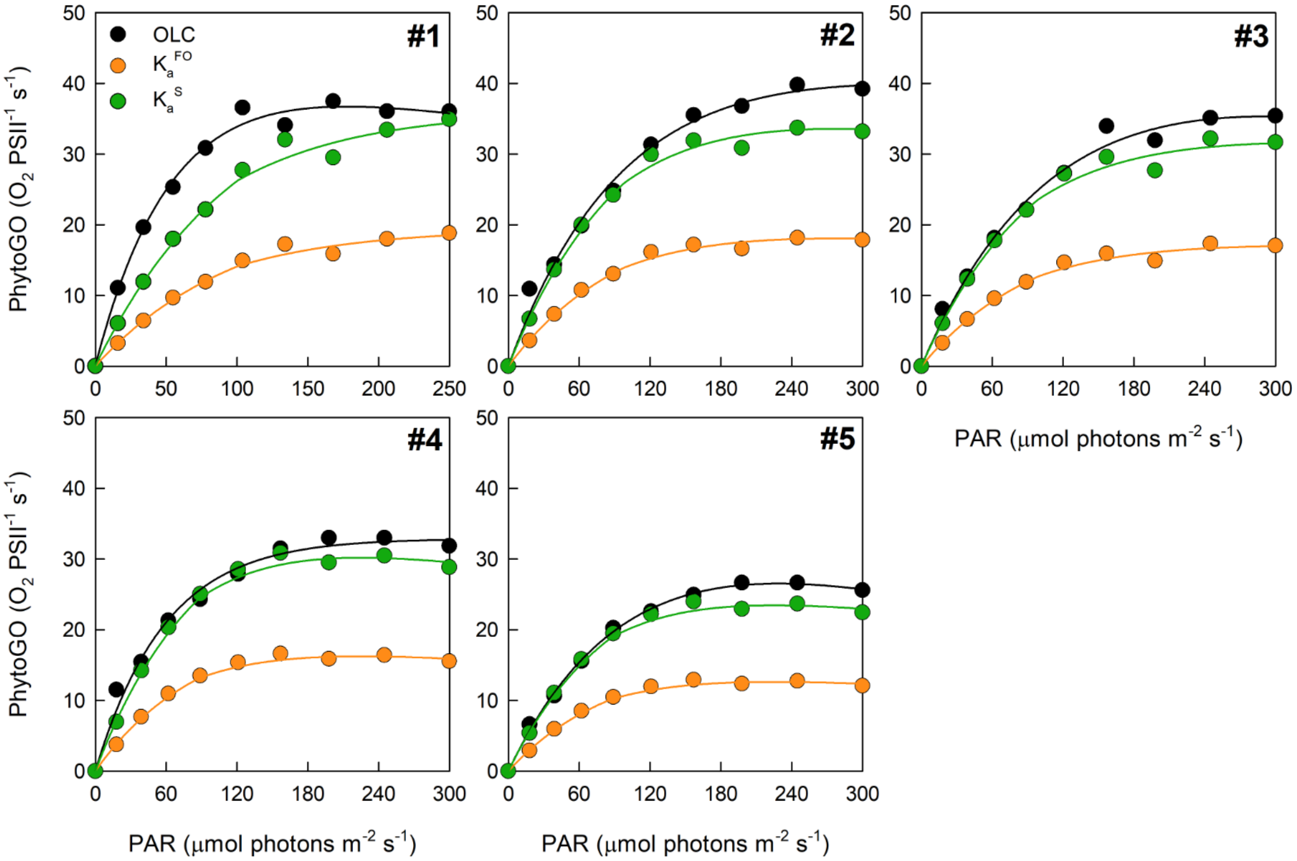


**Supplementary Figure 8.** The replicate simultaneous oxygen light curve (OLC) and fluorescence light curve (FLC) measurements made on *T. punctigera*. The OLC and FLC measurements were made on cultures acclimated to ambient temperature (~ 20 °C) and low-light (LL = 30 µmol photons $m^{-2}$ $s^{-1}$). FLC data was standardized to equivalent units of $O_2$, with both OLC and FLC data normalized to a derived concentration of PSII reaction centers (RCII) ($O_2$ RCII $s^{-1}$). FLC data were calculated using $K_a^{FO}$ = 11,800 $m^{-1}$ or the sample-specific $K_a^S$ value of 21,927 $m^{-1}$. The solid lines represent the PE curve fits. Each panel represents a biological replicate.

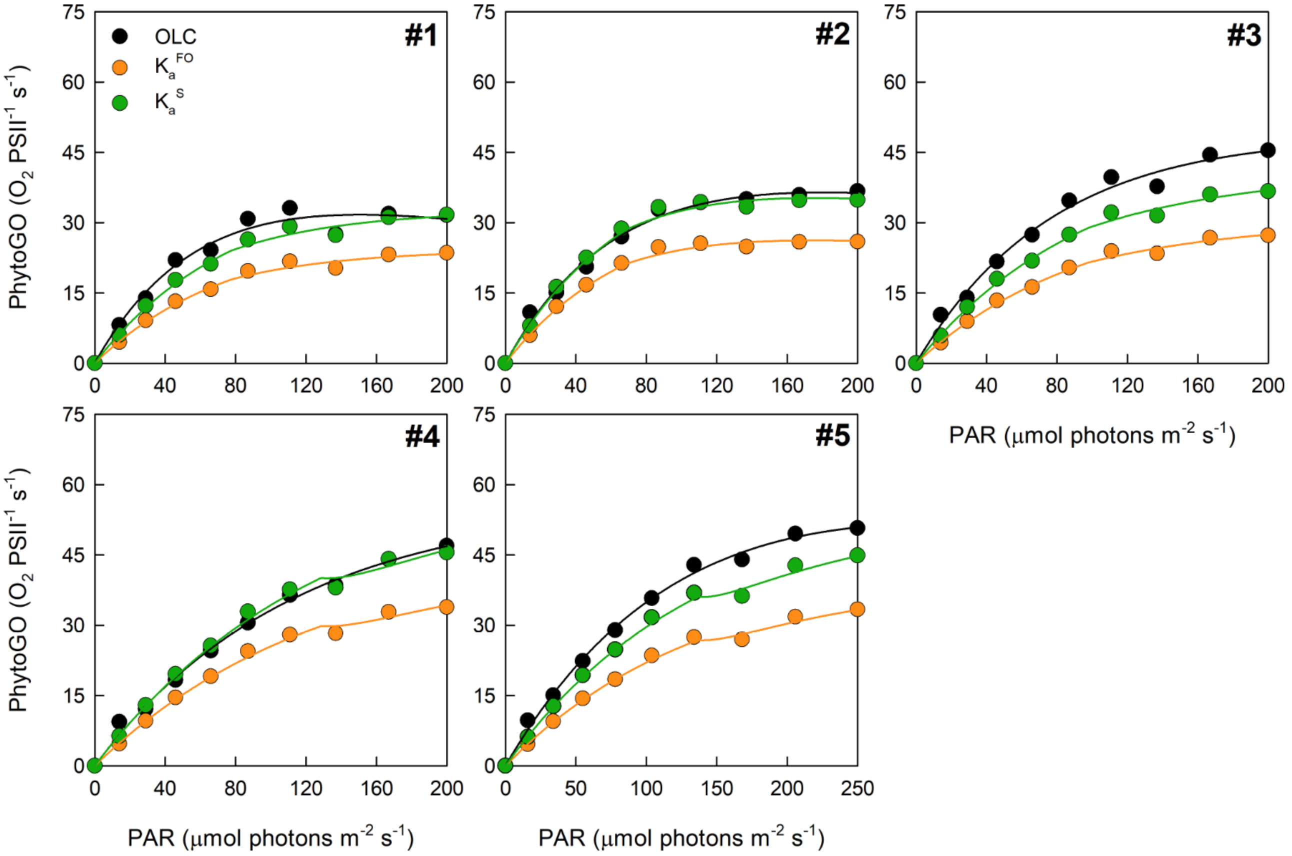


**Supplementary Figure 9.** The replicate simultaneous oxygen light curve (OLC) and fluorescence light curve (FLC) measurements made on *T. weissflogii*. The OLC and FLC measurements were made on cultures acclimated to ambient temperature (~ 20 °C) and low-light (LL = 30 µmol photons $m^{-2}$ $s^{-1}$). FLC data was standardized to equivalent units of $O_2$, with both OLC and FLC data normalized to a derived concentration of PSII reaction centers (RCII) ($O_2$ RCII $s^{-1}$). FLC data were calculated using $K_a^{FO}$ = 11,800 $m^{-1}$ or the sample-specific $K_a^S$ value of 15,868 $m^{-1}$. The solid lines represent the PE curve fits. Each panel represents a biological replicate.

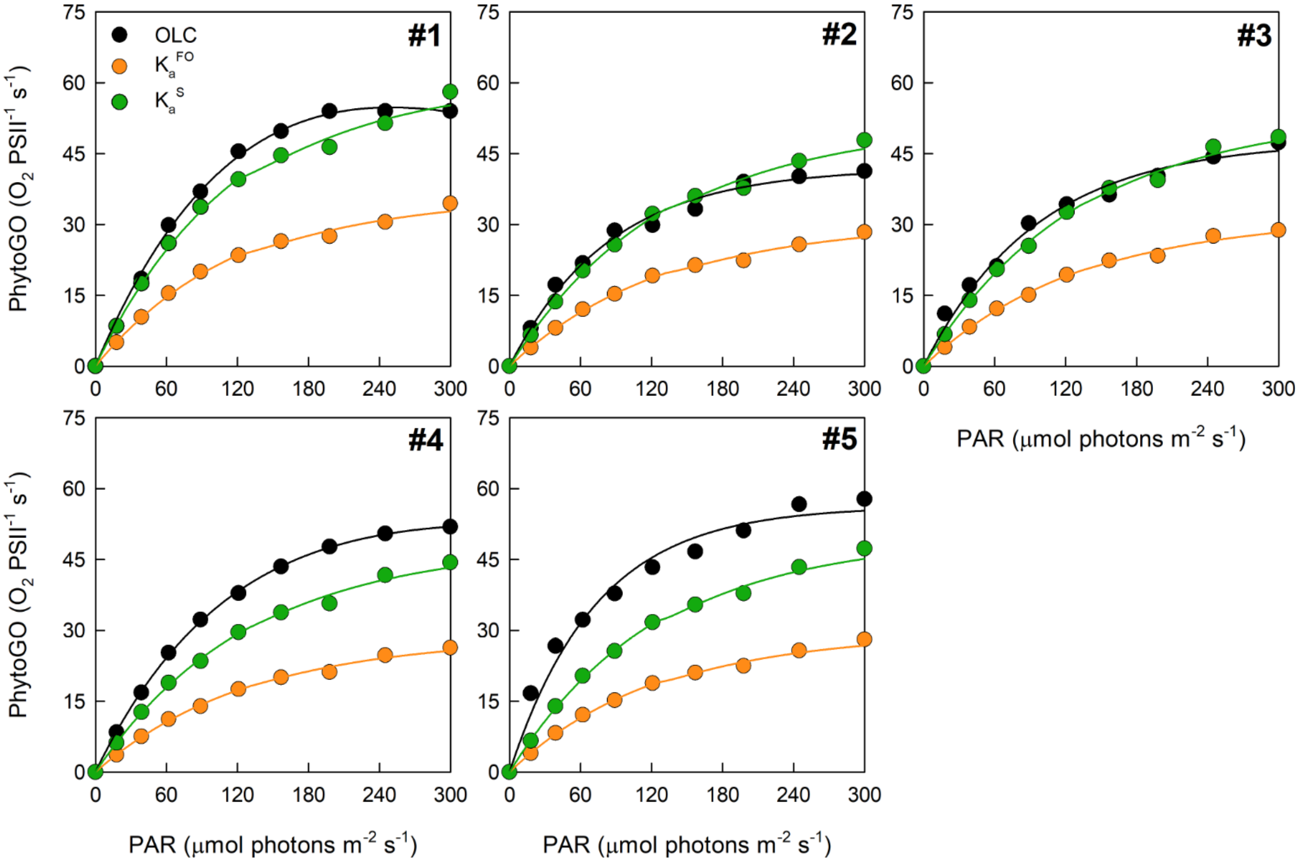


**Supplementary Figure 10.** The replicate simultaneous oxygen light curve (OLC) and fluorescence light curve (FLC) measurements made on *C. pelagicus*. The OLC and FLC measurements were made on cultures acclimated to ambient temperature (~ 20 °C) and low-light (LL = 30 µmol photons $m^{-2}$ $s^{-1}$). FLC data was standardized to equivalent units of $O_2$, with both OLC and FLC data normalized to a derived concentration of PSII reaction centers (RCII) ($O_2$ RCII $s^{-1}$). FLC data were calculated using $K_a^{FO}$ = 11,800 $m^{-1}$ or the sample-specific $K_a^{S}$ value of 19,898 $m^{-1}$. The solid lines represent the PE curve fits. Each panel represents a biological replicate.

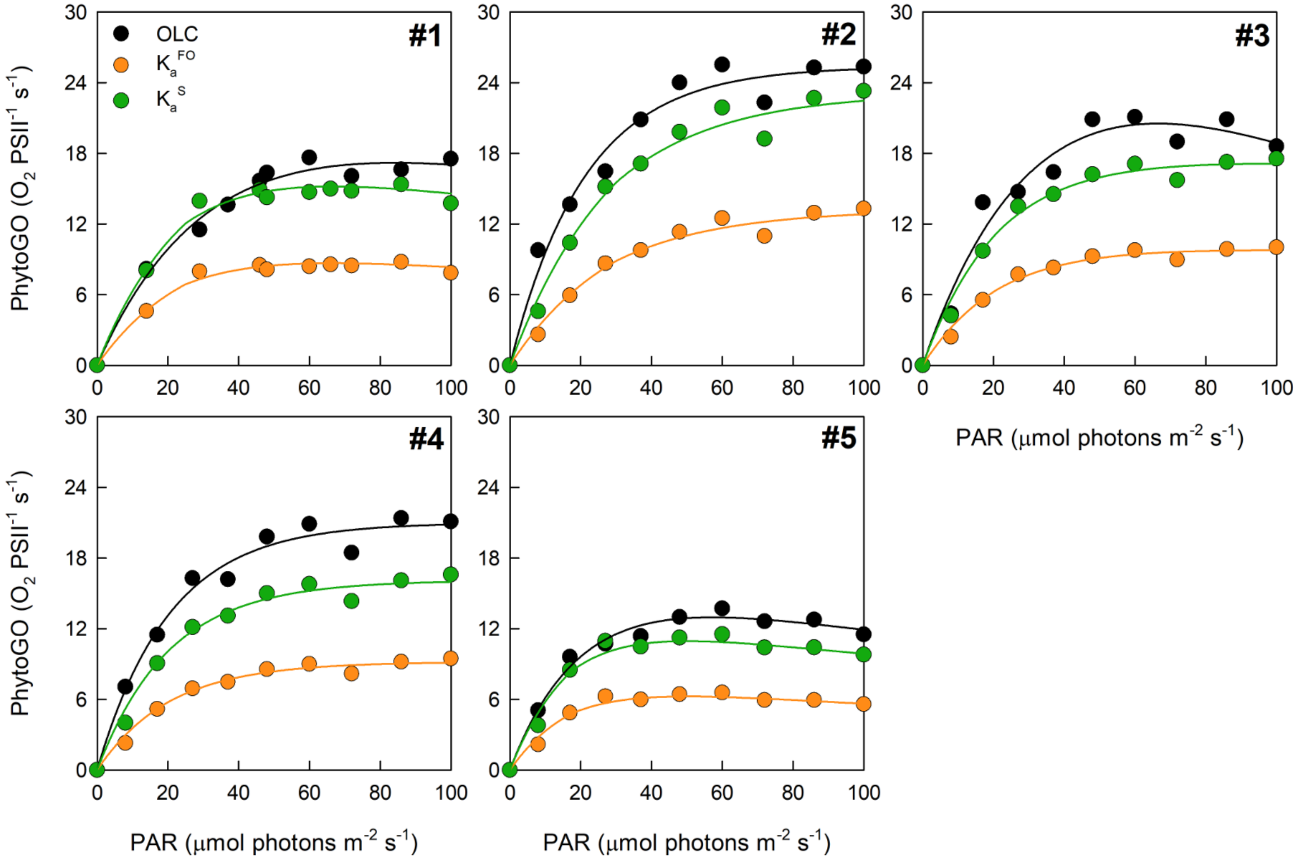


**Supplementary Figure 11.** The replicate simultaneous oxygen light curve (OLC) and fluorescence light curve (FLC) measurements made on *E. huxleyi*. The OLC and FLC measurements were made on cultures acclimated to ambient temperature (~ 20 °C) and low-light (LL = 30 µmol photons $m^{-2}$ $s^{-1}$). FLC data was standardized to equivalent units of $O_2$, with both OLC and FLC data normalized to a derived concentration of PSII reaction centers (RCII) ($O_2$ RCII $s^{-1}$). FLC data were calculated using $K_a^{FO}$ = 11,800 $m^{-1}$ or the sample-specific $K_a^{S}$ value of 20,677 $m^{-1}$. The solid lines represent the PE curve fits. Each panel represents a biological replicate.

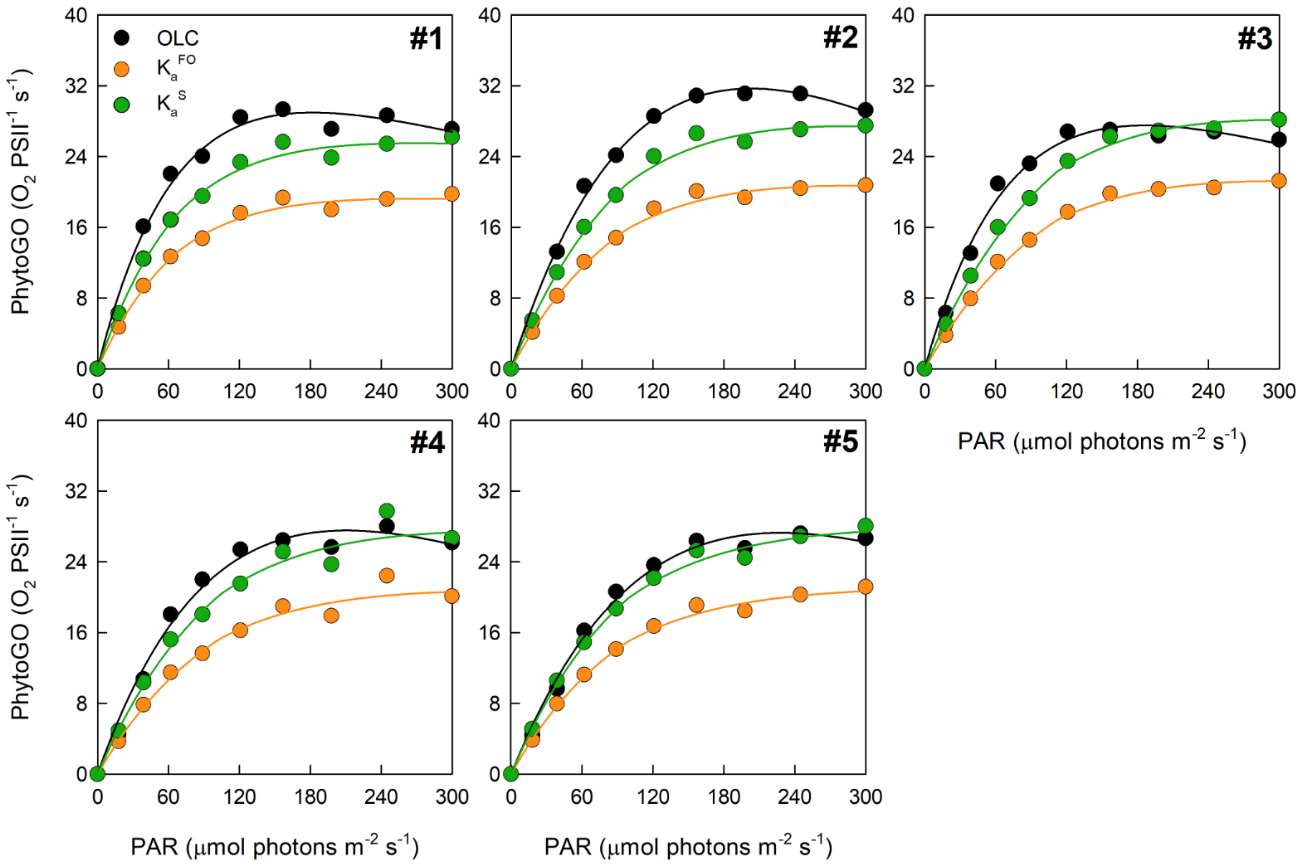


**Supplementary Figure 12.** The replicate simultaneous oxygen light curve (OLC) and fluorescence light curve (FLC) measurements made on *I. galbana*. The OLC and FLC measurements were made on cultures acclimated to ambient temperature (~ 20 °C) and low-light (LL = 30 μmol photons $m^{-2}$ $s^{-1}$). FLC data was standardized to equivalent units of $O_2$, with both OLC and FLC data normalized to a derived concentration of PSII reaction centers (RCII) ($O_2$ RCII $s^{-1}$). FLC data were calculated using $K_a^{FO}$ = 11,800 $m^{-1}$ or the sample-specific $K_a^S$ value of 15,644 $m^{-1}$. The solid lines represent the PE curve fits. Each panel represents a biological replicate.
